# Control of neurotransmitter release and presynaptic plasticity by re-orientation of membrane-bound Munc13-1

**DOI:** 10.1101/2021.07.29.454356

**Authors:** Marcial Camacho, Bradley Quade, Thorsten Trimbuch, Junjie Xu, Levent Sari, Josep Rizo, Christian Rosenmund

**Author notes:** These authors contributed equally: Bradley Quade, Marcial Camacho.

## Abstract

Munc13-1 plays a central role in neurotransmitter release through its conserved C-terminal region, which includes a diacyglycerol (DAG)-binding C_1_ domain, a Ca^2+^/PIP_2_-binding C_2_B domain, a MUN domain and a C_2_C domain. Munc13-1 was proposed to bridge synaptic vesicles to the plasma membrane in two different orientations mediated by distinct interactions of the C_1_C_2_B region with the plasma membrane: i) one involving a polybasic face that yields a perpendicular orientation of Munc13-1 and hinders release; and ii) another involving the DAG-Ca^2+^-PIP_2_-binding face that induces a slanted orientation and facilitates release. Here we have tested this model and investigated the role of the C_1_C_2_B region in neurotransmitter release. We find that K603E or R769E point mutations in the polybasic face severely impair synaptic vesicle priming in primary murine hippocampal cultures, and Ca^2+^-independent liposome bridging and fusion in in vitro reconstitution assays. A K720E mutation in the polybasic face and a K706E mutation in the C_2_B domain Ca^2+^-binding loops have milder effects in reconstitution assays and do not affect vesicle priming, but enhance or impair Ca^2+^-evoked release, respectively. The phenotypes caused by combining these mutations are dominated by the K603E and R769E mutations. Our results show that the C_1_-C_2_B region of Munc13-1 plays a central role in vesicle priming and support the notion that re-orientation of Munc13-1 controls neurotransmitter release and short-term presynaptic plasticity.

---

The release of neurotransmitters by Ca^2+^-evoked synaptic vesicle exocytosis is a central event in neuronal communication. This process involves a series of steps that include tethering of synaptic vesicles (SVs) to specialized areas of the plasma membrane called active zones, priming of the vesicles to a release-ready state and Ca^2+^-triggered fusion of the vesicle and plasma membranes (Sudhof, 2013). Neurotransmitter release does not merely constitute a means to transmit signals between neurons. The efficiency of release is regulated by a wide variety of mechanisms during presynaptic plasticity processes that shape the properties of neural networks and underlie multiple forms of information processing in the brain (Regehr, 2012). Hence, presynaptic terminals can be viewed as minimal computational units of the brain, and understanding the mechanisms that modulate neurotransmitter release is crucial to understand brain function.

The sophisticated protein machinery that controls neurotransmitter release has been extensively characterized (Brunger et al., 2018; Rizo, 2018), yielding defined models for the functions of the central components of this machinery and allowing reconstitution of fundamental features of synaptic exocytosis in liposome fusion assays that reproduce the critical functional importance of each one of these components (Ma et al., 2013; Stepien and Rizo, 2021). The SNAP receptor (SNARE) proteins syntaxin-1, SNAP-25 and synaptobrevin play a key role in membrane fusion by forming a tight SNARE complex that consists of a four-helix bundle and brings the vesicle and plasma membranes together (Hanson et al., 1997; Poirier et al., 1998; Sollner et al., 1993; Sutton et al., 1998). N-ethyl maleimide sensitive factor (NSF) and soluble NSF attachment proteins (SNAPs) disassemble the cis-SNARE complexes that result after fusion, recycling the SNAREs (Mayer et al., 1996; Sollner et al., 1993). NSF and SNAPs also dissociate trans-SNARE complexes and other four-helix bundles formed by the SNAREs (Choi et al., 2018; Ma et al., 2013; Prinslow et al., 2019; Yavuz et al., 2018), thus hindering fusion mediated by the SNAREs alone but at the same time ensuring that release occurs through a highly regulated fusion pathway that requires Munc18-1 and Munc13s (Park et al., 2017; Stepien et al., 2019), and that ensures proper parallel assembly of the SNARE four-helix bundle (Lai et al., 2017). Munc18-1 and Munc13s are essential for neurotransmitter release (Augustin et al., 1999; Richmond et al., 1999; Varoqueaux et al., 2002; Verhage et al., 2000) and organize trans-SNARE complex assembly via an NSF-SNAP-resistant pathway that starts with Munc18-1 bound to a self-inhibited ‘closed’ conformation of syntaxin-1 (Dulubova et al., 1999; Misura et al., 2000). Munc18-1 also binds to synaptobrevin, forming a template to assemble the SNARE complex (Baker et al., 2015; Jiao et al., 2018; Parisotto et al., 2014; Sitarska et al., 2017) while Munc13-1 bridges the vesicle and plasma membranes (Liu et al., 2016; Quade et al., 2019) and helps to open syntaxin-1 (Ma et al., 2011; Yang et al., 2015). This pathway ensures that release strictly requires Munc13s (Park et al., 2017), thus enabling the multiple forms of presynaptic plasticity that depend on these proteins.

Munc13s act as master regulators of release through their multidomain architecture. Munc13-1, the major isoform in brain, includes a variable N-terminal region and a C-terminal region that is conserved in all Munc13 isoforms (Fig. 1A). The N-terminal region contains a calmodulin-binding region involved in Ca^2+^-dependent short-term plasticity (Junge et al., 2004) and a C_2_A domain that forms a homodimer as well as a heterodimer with αRIMs, coupling RIM-dependent forms of plasticity to the priming machinery (Betz et al., 2001; Camacho et al., 2017; Deng et al., 2011; Dulubova et al., 2005; Lu et al., 2006). The conserved C-terminal region of Munc13-1 includes: i) the C_1_ domain, which mediates diacylglycerol (DAG)/phorbol ester-dependent potentiation of release (Basu et al., 2007; Rhee et al., 2002); ii) the C_2_B domain, which binds Ca^2+^ and phosphatidylinositol 4,5-bisphosphate (PIP_2_) and is involved in Ca^2+^- dependent short-term plasticity (Shin et al., 2010); iii) the MUN domain, which facilitates syntaxin-1 opening (Ma et al., 2011; Magdziarek et al., 2020); and iv) the C_2_C domain, which binds weakly to membranes in a Ca^2+^-independent manner (Quade et al., 2019). The crystal structure of a Munc13-1 fragment containing the C_1_, C_2_B and MUN domains (Xu et al., 2017) revealed that the C_1_ and C_2_B domains pack at one end of the highly elongated MUN domain, with their DAG- and Ca^2+^/PIP_2_-binding sites pointing to the same direction (Fig. 1A). The C_2_C domain [illustrated by a homology model of the C_2_C domain (Quade et al., 2019) in Fig. 1A] is expected to emerge on the opposite end of the MUN domain. This architecture enables the membrane bridging activity of Munc13-1, which led to a model whereby the C_2_C domain binds to the vesicle membrane while the C_1_-C_2_B region binds to the plasma membrane (Liu et al., 2016; Quade et al., 2019; Xu et al., 2017). This model was supported by the finding that mutations in predicted membrane-binding residues of the C_2_C domain dramatically impaired the liposome clustering ability of a fragment spanning the Munc13-1 C_1_, C_2_B, MUN and C_2_C domains (C_1_C_2_BMUNC_2_C) in vitro, as well as vesicle docking and neurotransmitter release in neurons (Quade et al., 2019).

**Figure 1.**
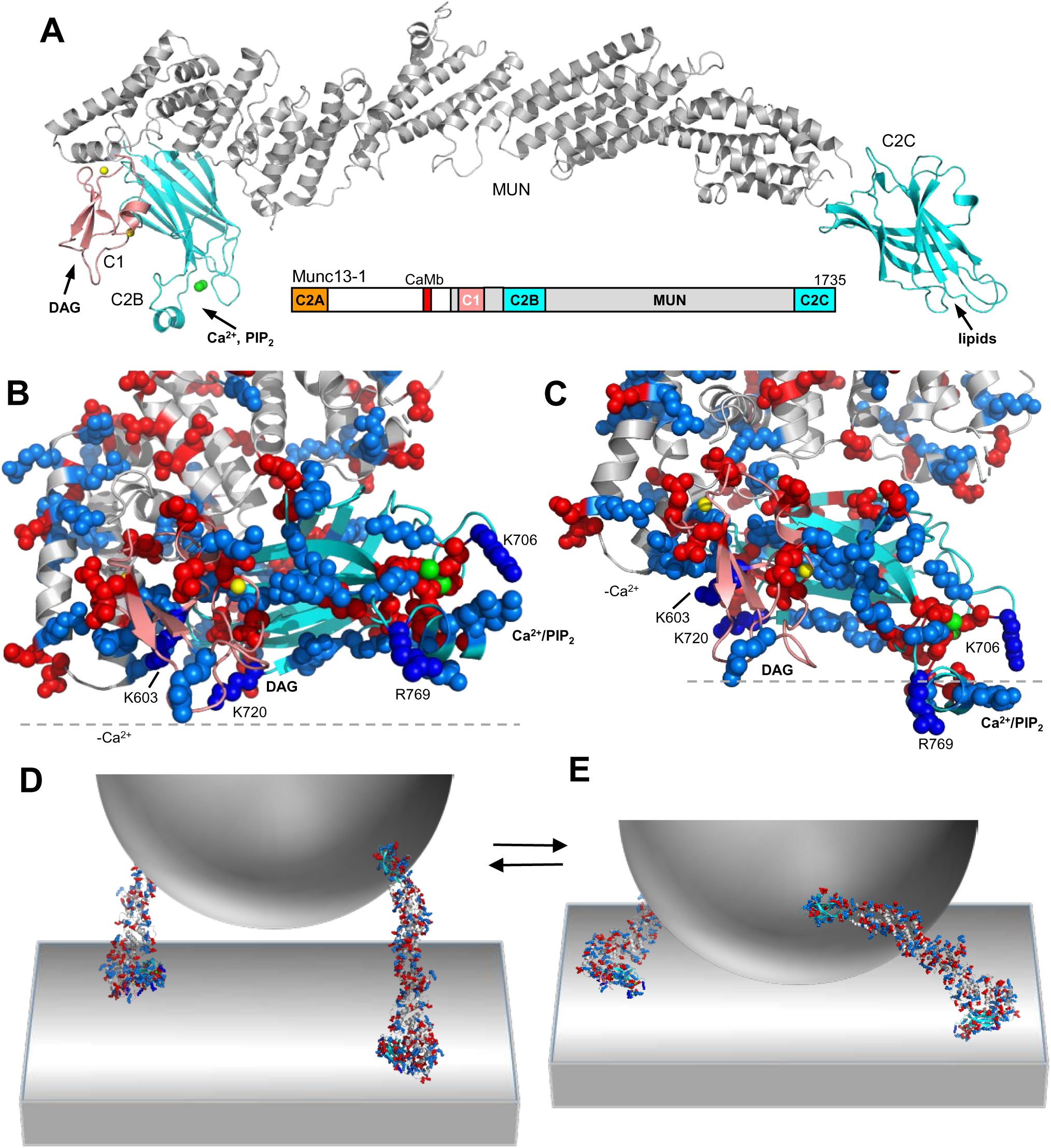
Model of Munc13-1 function with two different orientations bridging the membranes. (**A**) Domain diagram of Munc13-1 (CaMb, calmodulin binding domain) and ribbon diagrams illustrating a model of the structure of the Munc13-1 C_1_C_2_BMUNC_2_C fragment. The grey ribbon diagram is based on the crystal structure of the Munc13-1 C_1_C_2_BMUN fragment (Xu et al., 2017) (PDB accession number 5UE8) but, because some residues in loops of the C_1_ and C_2_B domains of this structure are missing, these domains were replaced with the NMR structure of the C_1_ domain (Shen et al., 2005) (salmon, PDB accession number 1Y8F) and the crystal structure of the Ca^2+^-bound C_2_B domain (Shin et al., 2010) (cyan, PDB accession number 3KWU). The cyan ribbon diagram on the right is a model of the C_2_C domain built by homology with the RIM1α C_2_B domain (Guan et al., 2007) (PDB accession code 2Q3X). (**B,C**) Close up views of the structure of the model of the Munc13-1 C_1_C_2_BMUN fragment shown in **A**. The dashed lines indicate the positions of planes parallel to the plasma membrane in the orientations predicted when binding is mediated by the polybasic face (**B**) or the DAG/Ca^2+^/PIP_2_-binding face (**C**) of the C_1_-C_2_B region. The Zn^2+^ ions bound to the C_1_ domain and the Ca^2+^ ions bound to the C_2_B domain are shown as yellow or green spheres, respectively. Basic residues are shown as blue spheres and acidic residues as red spheres. The positions of the residues that were mutated in this study and the approximate locations of the DAG- and Ca^2+^/PIP_2_-binding sites are indicated. (**D,E**) Three-dimensional models that illustrate the notion that Munc13-1 C_1_C_2_BMUNC_2_C can bridge the synaptic vesicle and plasma membranes in two different orientations. The models include a vesicle (half sphere), the plasma membrane (flat surface) and ribbon diagrams representing the modelled structure of the C_1_C_2_BMUNC_2_C fragment shown in **A** in the approximately perpendicular (**D**) and slanted (**E**) orientations with respect to the plasma membrane yielded by the MD simulations performed in the absence and presence of Ca^2+^, respectively. The model postulates that perpendicular orientation allows partial but not full assembly of the SNARE complex, whereas the slanted orientation facilitates full assembly. The two states are proposed to exist in an equilibrium that is shifted to the right by Ca^2+^ and DAG.

The advances made in understanding the core of the release machinery provide a framework to elucidate the molecular mechanisms that underlie Munc13-1-dependent presynaptic plasticity. A model of how increases in DAG and intracellular Ca^2+^ levels during repetitive stimulation facilitate vesicle priming and neurotransmitter release emerged from the crystal structure of a Munc13-1 C_1_C_2_BMUN fragment and the realization that the C_1_ and C_2_B domains could bind to the plasma membrane through their DAG- and Ca^2+^/PIP_2_-binding sites in a Ca^2+^-dependent manner, and through a polybasic region that partially overlaps with these sites in the absence of Ca^2+^ (referred to below as the polybasic face; see Fig. 1B,C) (Xu et al., 2017). These observations, together with liposome clustering and liposome fusion data, suggested that Munc13-1 can bridge a vesicle to the plasma membrane in two orientations (Liu et al., 2016; Quade et al., 2019; Xu et al., 2017): a close to perpendicular orientation that involves the polybasic face of the C_1_-C_2_B region is favored in the absence of Ca^2+^ and facilitates partial assembly of SNARE complexes but hinders progress toward membrane fusion; and a more slanted orientation that also involves the C_1_-C_2_B region is favored by Ca^2+^ and DAG, and stimulates further SNARE complex assembly and fusion (Fig. 1D,E). Interestingly, a large amount of experimental data on short-term presynaptic plasticity accumulated over the years can be explained by a related model whereby there is a dynamic equilibrium between two primed states that involve different orientations of Munc13-1 and different extents of SNARE complex assembly, and the equilibrium can be shifted by Ca^2+^ and DAG (Neher and Brose, 2018). The proposed dual inhibitory and stimulatory roles of the membrane bridging activity of Munc13-1 are also consistent with the effects of point mutations that disrupt interactions between the various Munc13-1 domains (Xu et al., 2017), with the finding that a H567K mutation that unfolds the C_1_ domain of Munc13-1 impairs vesicle priming but increases the vesicular release probability (Basu et al., 2007; Rhee et al., 2002), and with the observation that deletion of the C_1_ or C_2_B domain of unc-13 enhances neurotransmitter release in *C. elegans*, yet deletion of both domains strongly hinders release (Michelassi et al., 2017). Nevertheless, the validity of this two state model and the functional importance of the polybasic face formed by the C_1_ and C_2_B domains remain to be demonstrated.

The study presented here was designed to test this model and investigate the functional consequences of mutations in predicted membrane-binding residues of the Munc13-1 C_1_-C_2_B region using a combination of molecular dynamics (MD) simulations, electrophysiological experiments in neurons and biochemical and reconstitution assays in vitro. We find that K603E or R769E single point mutations in the C_1_C_2_B polybasic face of Munc13-1 disrupt Ca^2+^-independent liposome binding and bridging, as well as stimulation of liposome fusion in reconstitution assays, and severely impair synaptic vesicle priming in mice autaptic cultures. Conversely, a K720E mutation in the polybasic face and a K706E mutation in the C_2_B domain Ca^2+^-binding loops, which have milder effects on liposome binding, bridging and fusion, do not affect vesicle priming; however, Ca^2+^-evoked release and the release probability are enhanced by the K720E mutation and impaired by K706E. The phenotypes caused by combining these mutations are dominated by the K603E and R769E mutations. These results strongly support the notion that binding of the C_1_-C_2_B region of Munc13-1 to the plasma membrane is critical for synaptic vesicle priming and that a switch from a perpendicular to a slanted orientation of Munc13-1 controls neurotransmitter release and short-term presynaptic plasticity.

## Results

### Models of Munc13-1 C_1_-C_2_B-membrane interactions in the absence and presence of Ca^2+^

The notion that the C_1_-C_2_B region of Munc13-1 can bind to membranes via different surfaces in the absence and presence of Ca^2+^ emerged from analysis of the distribution of charged residues in these surfaces (Xu et al., 2017) (Fig. 1B,C). To derive models of these putative membrane-binding modes, we performed MD simulations using a model of a fragment spanning the C_1_, C_2_B and MUN domains of Munc13-1 (C_1_C_2_BMUN) based on its crystal structure (Xu et al., 2017), and a square membrane with a lipid composition that mimics that of the plasma membrane (Chan et al., 2012). In one simulation, we placed the Ca^2+^-free C_1_C_2_BMUN model above the cytoplasmic leaflet in an approximately perpendicular orientation with the polybasic face close to the membrane, and ran a simulation of 100 ns. The orientation of C_1_C_2_BMUN became even more perpendicular in the first 10 ns and during the remaining 90 ns oscillated around an almost completely perpendicular orientation such as that observed at the end of the simulation (Figure 1-Figure supplement 1A,B). The C_1_-C_2_B region quickly established extensive interactions with the membrane, including multiple salt bridges between basic residues and the phospholipid head groups (Figure 1-Figure supplement 2A).

We carried a second simulation in which we included two Ca^2+^ ions bound to the corresponding binding sites of the C_2_B domain (Shin et al., 2010) and C_1_C_2_BMUN was placed in a more slanted orientation such that the DAG- and Ca^2+^/PIP_2_-binding sites of the C_1_ and C_2_B domains, respectively, were close to the membrane. A range of slanted orientations of C_1_C_2_BMUN were visited during the 86 ns simulation, oscillating during the last 70 ns around the orientation observed in the last pose, which was even more slanted than the initial orientation (Figure 1-Figure supplement 1C,D). During the simulation, the expected DAG and PIP_2_ binding regions of the C_1_ and C_2_B domains became intimately bound to the membrane through numerous interactions involving hydrophobic and basic residues of both domains (Figure 1-Figure supplement 2B). Interestingly, the unique amphipathic α-helix present in one of the Ca^2+^-binding loops of the C_2_B domain (Shin et al., 2010) inserted partially into the membrane through its hydrophobic side chains while its basic residues interacted with the phospholipid head groups. It is noteworthy that this helix also interacts with the membrane in the perpendicular orientation adopted in the Ca^2+^-free simulation, which allows a more extensive membrane-interacting surface involving the sides of the C_1_ and C_2_B domains, as well as the linker between them (Figure 1-Figure supplements 1B and 2A). In the slanted, Ca^2+^-bound orientation, the interaction surface is smaller because it involves only the tips of the C_1_ and C_2_B domain, but hydrophobic groups of the C_2_B domain helix and the C_1_ domain insert into the membrane, and a PIP_2_ head group located between the Ca^2+^-binding loops is close to the Ca^2+^ ions (Figure 1-Figure supplements 1D and 2B). Since phospholipid head groups are known to complete the coordination spheres of Ca^2+^ ions bound to C_2_ domains, dramatically increasing their Ca^2+^ affinity (Verdaguer et al., 1999; Zhang et al., 1998), it seems likely that phospholipid-Ca^2+^ coordination and the insertion of hydrophobic groups into the membrane provide the driving force for this binding mode and a Ca^2+^-induced change in orientation of Munc13-1 with respect to the membrane.

We would like to emphasize that these simulations were biased by the initial orientations and a much more thorough analysis will be required to explore the possible binding modes of the Munc13-1 C_1_- C_2_B region with the plasma membrane. However, these short simulations were dynamically very stable around the two distinct basins, with and without Ca^2+^, supporting the notion that Munc13-1 can adopt two dramatically different orientations with respect to the plasma membrane. The simulations yielded chemically reasonable binding modes and help to visualize how changes in the orientation of Munc13-1 can drastically alter the distance between a vesicle and the plasma membrane (Fig. 1D,E). Note also that, although we postulate that Ca^2+^ favors the switch from the perpendicular to the slanted orientation, both binding modes are likely possible in the absence and presence of Ca^2+^, and other factors such a more complete assembly of the SNARE complex may induce the switch to the slanted orientation even in the absence of Ca^2+^. The observed binding modes provide useful frameworks for the design of mutations to disrupt Munc13-1-membrane interactions involving the C_1_C_2_B region. Using these models, we designed the following mutations to investigate the functional importance of the distinct faces of the C_1_-C_2_B region and test the hypothesis that this region mediates binding of Munc13-1 to membranes in two different orientations that underlie in part its role in regulating neurotransmitter release: i) K603E and K720E mutations in the C_1_ and C_2_B domain, respectively, probed residues in the polybasic face that are far from the Ca^2+^-binding sites of the C_2_B domain and are predicted to interact with the lipids in the Ca^2+^- independent binding mode (Fig. 1B, Figure 1-Figure supplement 2A); ii) an R769E mutation probed the importance of a basic residue from one of the Ca^2+^-binding loops of the C_2_B domain that is involved in interactions with the lipids in both the Ca^2+^-independent and Ca^2+^-dependent binding modes yielded by the simulations (Fig. 1B,C, Figure 1-Figure supplement 2A,B); and iii) a K706E mutation in a residue from another Ca^2+^-binding loop of the C_2_B domain (loop 1) that is involved in lipid interactions in the Ca^2+^- dependent membrane binding mode but is far from the membrane in the Ca^2+^-independent mode (Fig. 1B,C, Figure 1-Figure supplement 2A,B). Note that a mutation of this residue to Trp in Munc13-2 led to a gain-of-function, increasing the Ca^2+^ sensitivity of neurotransmitter release (Shin et al., 2010).

### Differential disruption of neurotransmitter release by mutations in the C_1_C_2_B region of Munc13-1

To investigate the functional consequences of the K603E, K706E, K720E and R769E mutations in the C_1_C_2_B region of Munc13-1, we analyzed synaptic responses in autaptic hippocampal cultures from Munc13-1/2 double knockout (DKO) mice rescued with Munc13-1 carrying these point mutations individually. All expressed Munc13-1 mutants exhibited Munc13-1 fluorescence at VGLUT1-positive compartments, showing their presynaptic localization, and had similar expression levels as Munc13-1 WT (Figure 2-Figure supplement 1). We first analyzed their impact on SV priming as measured by depletion of the readily-release pool (RRP) of vesicles induced by hypertonic solution (Rosenmund and Stevens, 1996). We found that DKO neurons rescued with the Munc13-1 bearing the K603E mutation in the C_1_ domain or the R769E mutation in one of the C_2_B domain Ca^2+^-binding loops exhibited a drastically reduced RRP (Fig. 2A). However, no significant impact on RRP size was observed for the K720E mutation in the C_2_B domain, despite the proximity of K720 to K603 (Fig. 1B), or for the K706E mutation in another C_2_B domain Ca^2+^- binding loop (Fig. 2A).

**Figure 2:**
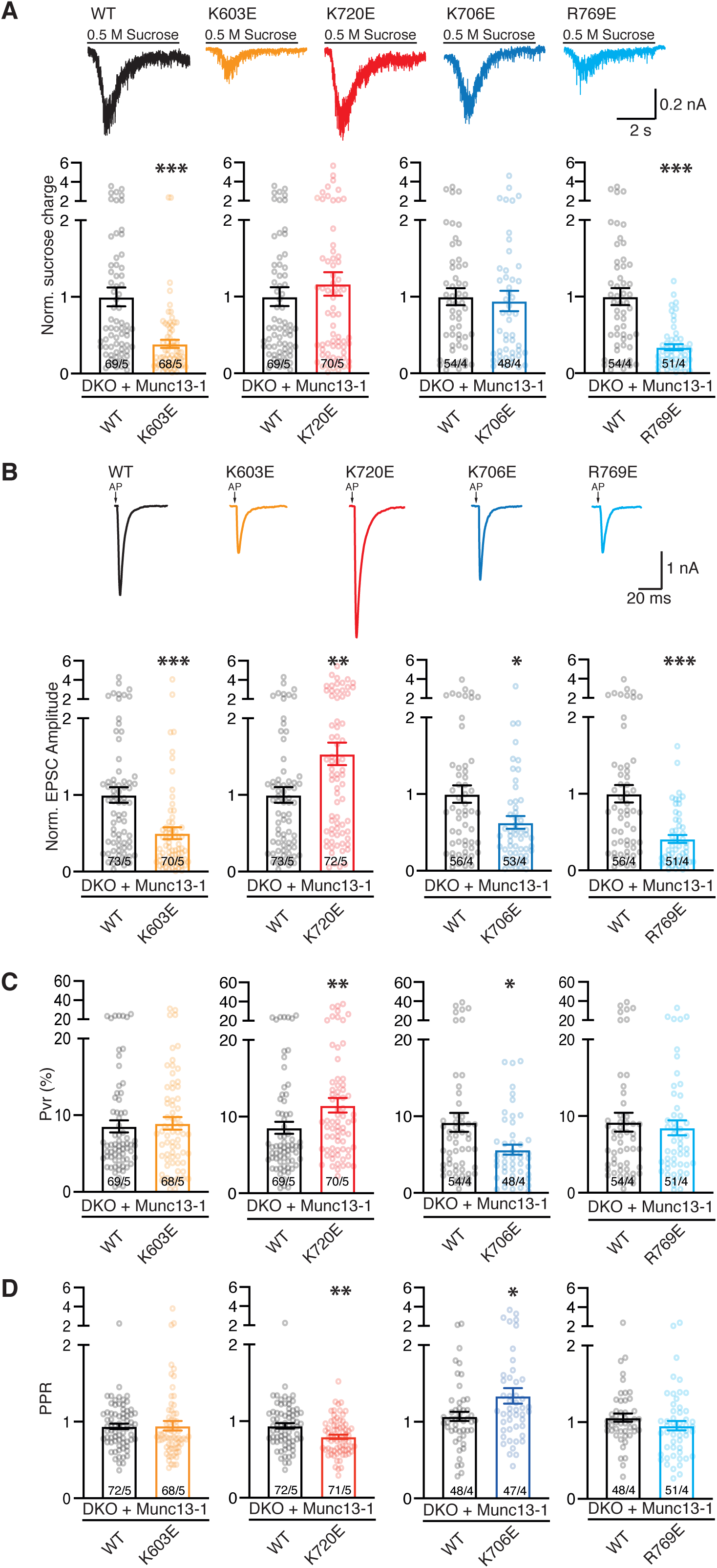
Evaluation of Munc13-1 functions using single point mutations within the C1-C2B region. (**A-D**) Electrophysiological parameters of autaptic hippocampal glutamatergic Munc13-1/2 DKO neurons expressing Munc13-1 WT (black) or Munc13-1 polybasic mutants: K603E (orange), K720E (red), K706E (blue) and R769 (light blue). (**A**) Representative sucrose-induced current traces and bar plots depicting the mean charge of the current evoked by the application of 500 mM hypertonic sucrose solution for 5s normalized to the corresponding Munc13-1 WT control. (**B**) Representative traces of AP-evoked EPSCs and bar plots showing the mean EPSC amplitudes normalized to the corresponding Munc13-1 WT. EPSC recordings were done at RT in 2mM Ca^2+^/4 mM Mg^2+^. Action potentials in traces were blanked for better illustration and substituted by arrows. (**C**) Bar plots depicting the average vesicular release probability (Pvr) calculated for each neuron. (**D**) Bar plots depicting average pair-pulse ratio (PPR) with a 25 ms interpulse interval. Circles in bar plot represent values per neuron. Numbers in bars correspond to the cell number/culture number. Significance and p-values were determined by comparison with the corresponding Munc13-1 WT using the non-parametric Mann-Whitney U test. * p <0.05; ** p <0.01; *** p <0.001.

We next examined the influence of the four mutations on Ca^2+^-evoked neurotransmitter release. The vesicular release probability (Pvr), the likelihood that an action potential causes the fusion of a primed and fusion competent vesicle, can be readily calculated by dividing the excitatory postsynaptic current (EPSC) and sucrose evoked charges (Reim et al., 2001; Rosenmund and Stevens, 1996). The K603E and R769E mutations decreased Ca^2+^-evoked release but did not alter the Pvr significantly (Fig. 2B,C). In contrast, the K706E mutation decreased both the EPSC amplitude and the Pvr, while the K720E mutation increased both evoked release and the Pvr (Fig. 2B,C). These effects on the efficiency of release observed for the K706E and K720E mutants were further confirmed by recordings with a pair pulse protocol consisting of two consecutive AP-induced EPSCs with an inter-stimulus interval of 25 ms (Fig. 2D). Thus, the K603E and R769E mutations did not alter the paired-pulsed ratio (PPR), but the K706E mutant exhibited an increased PPR, which is typical of synapses with low Pvr, whereas we observed a decreased PPR for the K720E mutant, as expected from its increased Pvr. We also assessed the impact of these mutations on spontaneous release by analyzing the frequency of miniature postsynaptic currents (mEPSCs). We observed a decrease in mEPSC frequency for K603E and R769E, the two mutants that exhibited impaired priming, whereas the K706E mutation did not affect spontaneous release and K720E, the mutant with enhanced Pvr, had increased mEPSC frequency (Fig. Figure 2-Figure supplement 2A).

Overall, these results show that two basic residues in the C_1_C_2_B region of Munc13-1, K603 and R769, play important roles in SV priming, whereas two other basic residues, K706 and K720, modulate the vesicular release probability and have opposite effects. To gain further insights into the roles of these residues, we analyzed neurotransmitter release in rescue experiments with Munc13-1 double mutants that combined the two point mutations in residues that are far from the C_2_B domain Ca^2+^-binding sites (K603E/K720E) or the two mutations in the C_2_B domain Ca^2+^-binding loops (K706E/R769E), as well as a quadruple mutant where the four basic residues were replaced (K603E/K720E/K706E/R769E). The K603E/K720E mutant exhibited impaired sucrose-induced, Ca^2+^-evoked and spontaneous release, without significant changes in Pvr or PPR (Fig. 3, Figure 2-Figure supplement 2B), similar to the effects observed for the single K603E mutant. Hence, the K603E mutation dominates the phenotypes exhibited by this double mutant and cancels the gain-of-function caused by the single K720E mutation. The double point mutant K706E/R769E within the C_2_B domain Ca^2+^-binding loops displayed stronger defects in the size of the RRP, the EPSC amplitude and the spontaneous release frequency than the K603E/K720E mutant (Fig. 3A,B, Figure 2-Figure supplement 2B), and appeared to have a tendency to lower Pvr than wild type (WT), but the difference did not reach statistical significance (Fig. 3C). Correspondingly, the PPR observed for the K706E/R769E mutant showed a tendency to facilitation that was not statistically significant. Finally, we observed that the K603E/K720E/K706E/R769E quadruple mutant exhibited the lowest number of responsive cells and that these were strongly deficient in priming, spontaneous release and, particularly, Ca^2+^-evoked release, while the Pvr and RRP observed for this mutant had similar non-statistically significant tendencies as the K706E/R769E mutant (Fig. 3, Figure 2-Figure supplement 2B). These results show that the mutations that impair priming, K603E and R769E, dominate the phenotypes of these double and quadruple mutants and the effects of the mutations that specifically affect evoked release, K706E and K720E, are at least partially masked by these dominant effects.

**Figure 3:**
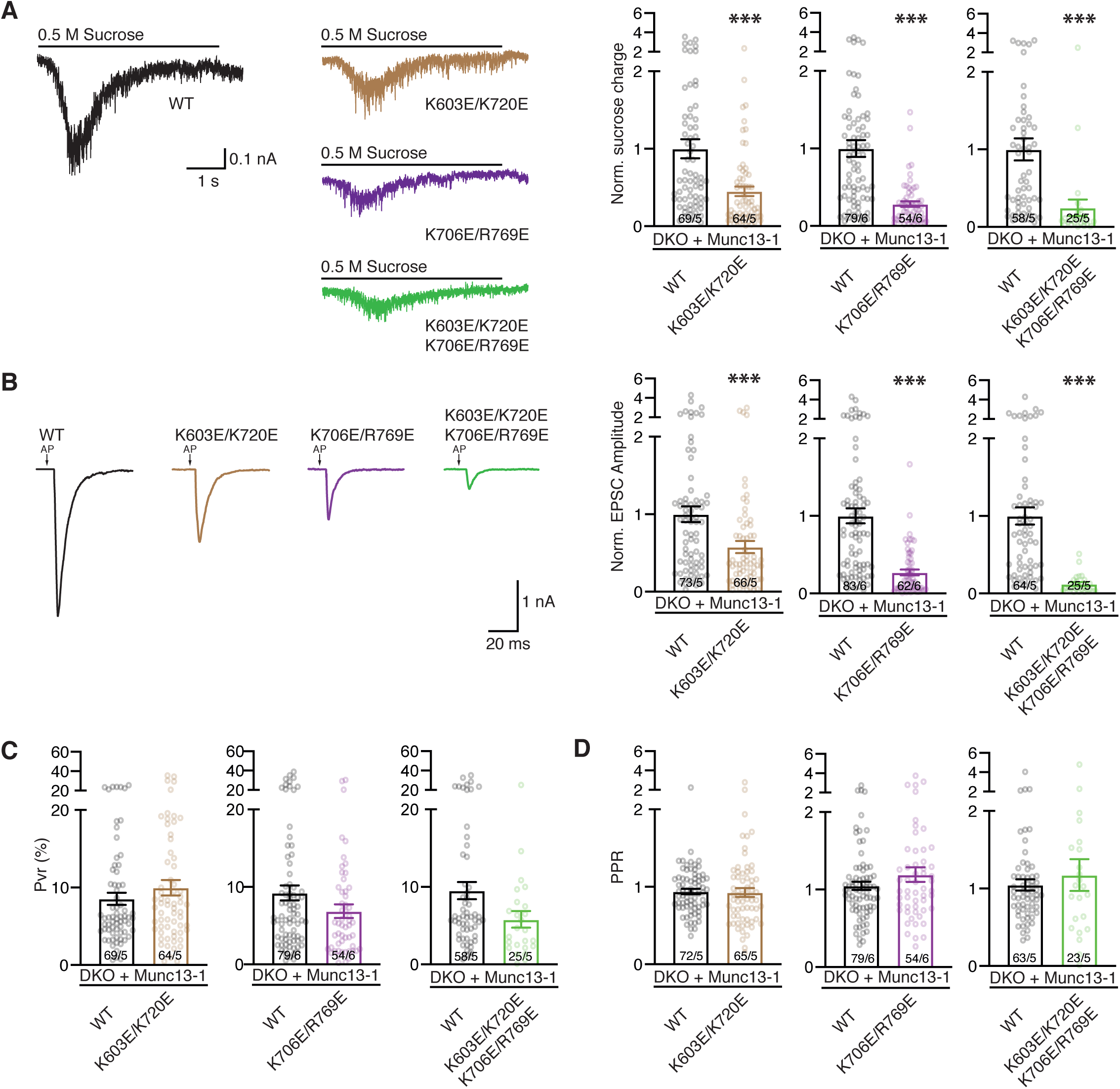
Phenotypic analyses of double and quadruple mutations within the Munc13-1 C1-C2B region. Representative sucrose-evoked synaptic current traces and bar plots of the charge currents normalized to the corresponding Munc13-1 WT control from autaptic hippocampal glutamatergic Munc13-1/2 DKO neurons rescued with Munc13-1 WT (black), Munc13-1 double point mutants, K603E/K720E (brown) or K706E/R769E (purple), or Munc13-1 quadruple point mutant K603E/K720E/K706E/R769E (green). (**B**) Representative AP-evoked EPSCs traces and bar plots of the normalized mean EPSC amplitudes from the groups above described. Action potentials in traces were blanked and substituted by arrows. (**C**) Bar plots showing the average vesicular release probability (Pvr) from the groups described in A and B. (**D**) Bar plots depicting average PPR at a frequency of 40 Hz. Circles in bar plot represent normalized values per neuron. Numbers in bars corresponded to the cell number/culture number. Significances and p-values were determined using the non-parametric Mann-Whitney U test. *P<0.05; ***P<0.001.

Overall, these data show that mutations in the C_1_C_2_B region of Munc13-1 can have effects on both SV priming and Ca^2+^-evoked release, and can lead to both loss-of-function and gain-of-function. These findings support the notion that the C_1_C_2_B region is involved in Ca^2+^-independent interactions through the polybasic face that are critical for priming and in additional interactions involving in part the C_2_B domain Ca^2+^-binding loops that affect the probability of evoked release, consistent with the two state model of Fig. 1D,E. The finding that mutations in two basic residues that are near each other (K603 and K720) lead to what appear to be opposite effects (loss- or gain-of-function) suggests that release is modulated by a delicate balance between inhibitory and stimulatory interactions involving the C_1_C_2_B region, which is also a key aspect of the model. It seems likely that at least some of the interactions involving the C_1_C_2_B polybasic face that mediate the initial bridging of the vesicle and plasma membranes by Munc13-1, leading to primed vesicles, need to be released for Munc13-1 to adopt the more slanted orientation that facilitates SNARE complex zippering and membrane fusion. The interplay between these two events may underlie the distinct effects of the K603E and K720E mutations, as both mutations can impair the initial bridging but facilitate the transition to the slanted orientation to different degrees (see discussion).

### Distinct effects of mutations in the C_1_C_2_B region of Munc13-1 on short-term plasticity

We next investigated how the mutations in basic residues of the C_1_C_2_B region affect synaptic responses during repetitive stimulation by applying 10 Hz stimulus trains on autaptic cultures of Munc13-1 DKO neurons rescued with WT and comparing them side-by-side with rescues by the Munc13-1 mutants. Analysis of these data is complicated by the fact that the observed changes in EPSC amplitudes during repetitive stimulation can be caused by various mechanisms, including depletion of the RRP, alterations in the efficiency of replenishment of the RRP and changes in the release probability. To help distinguishing between these mechanisms, we prepared plots of normalized responses, which inform on differences in the extent of depression or facilitation during the stimulus trains, and plots of absolute EPSC amplitudes, which can help to clarify the mechanisms related to use and re-fill of primed synaptic vesicles (Shin et al., 2010) (Fig. 4).

**Figure 4.**
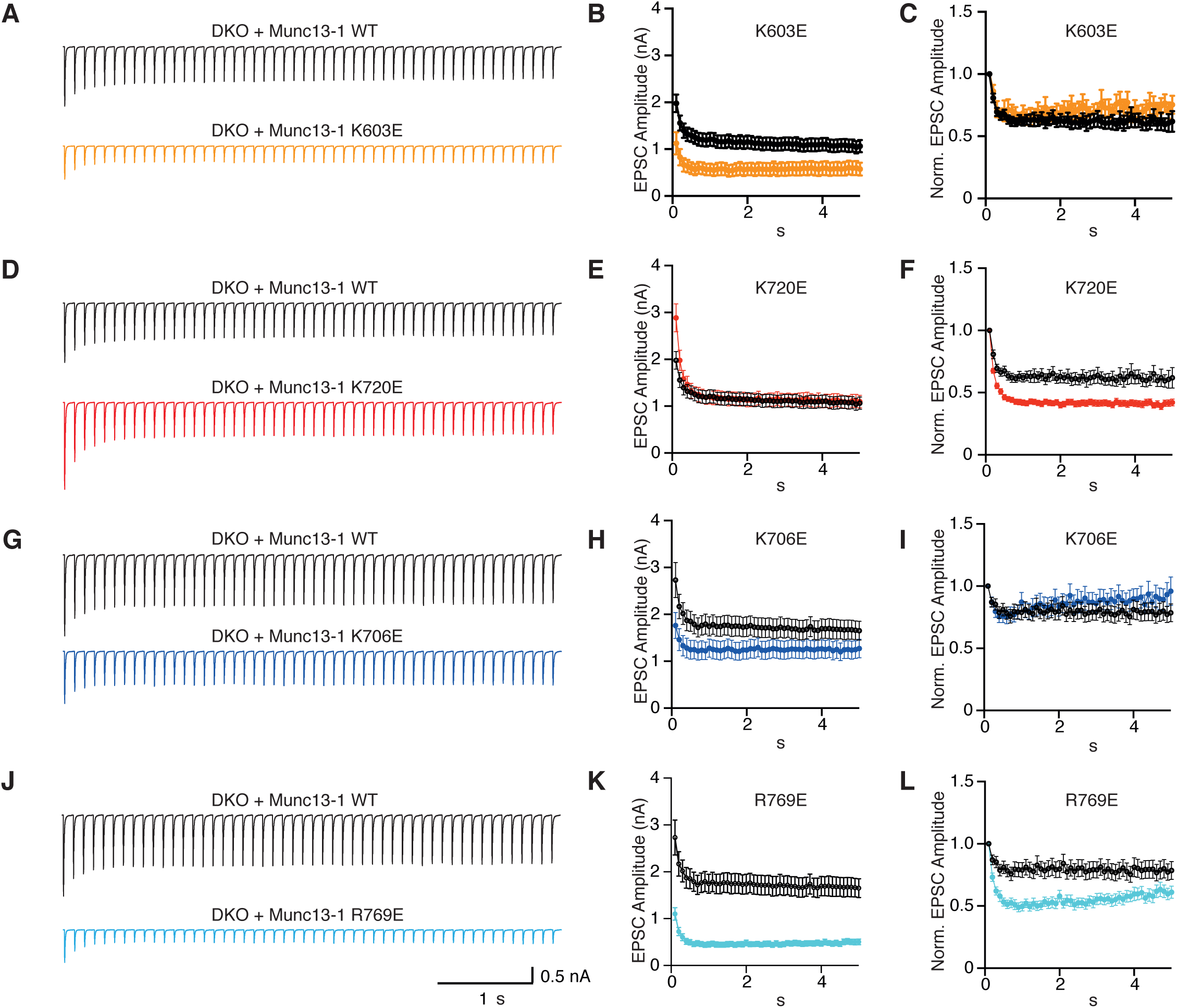
Short term plasticity behavior of single point mutations within the Munc13-1 C_1_-C_2_B region. (**A**) Average EPSC traces recorded during a 10 Hz train of 50 action potentials and summary graphs of absolute and normalized (**C**) EPSC amplitudes of autaptic hippocampal Munc13-1/2 DKO neurons expressing Munc13-1 WT (black) n=71/5 and Munc13-1 K603E (orange) n=67/5. Average 10 Hz train (**D**) and summary graphs of absolute (**E**) and normalized (**F**) EPSC amplitudes of autaptic hippocampal Munc13-1/2 DKO neurons rescued with Munc13-1 WT (black) n=71/5 and Munc13-1 K720E (red) n=67/5. Average 10 Hz train (**G**) and summary graphs of absolute (**H**) and normalized (**I**) EPSC amplitudes of autaptic hippocampal Munc13-1/2 DKO neurons rescued with Munc13-1 WT (black) n=54/4 and Munc13-1 K706E (blue) n=47/4. Average 10 Hz train (**J**) and summary graphs of absolute (**K**) and normalized (**L**) EPSC amplitudes of autaptic hippocampal Munc13-1/2 DKO neurons rescued with Munc13-1 WT (black) n=52/4 and Munc13-1 R769E (light blue) n=46/4. Action potentials artifacts were blanked.

As expected (Rosenmund et al., 2002), DKO neurons rescued with WT Munc13-1 displayed substantial depression in the beginning of the stimulus train that can be attributed to RRP depletion. The R769E single mutant exhibited stronger depression than WT Munc13-1 (Fig. 4L), which likely arises at least in part from impairment in the kinetics of RRP replenishment, as this mutant had a strong defect in priming but no deficit in Pvr (Fig. 2). However, the K603E mutant also had impaired priming and yet it exhibited similar depression to WT Munc13-1 in the beginning of the train, with a tendency to have higher EPSC amplitudes than WT as the train progressed (Fig. 4C). These findings suggest that the fusogenicity of primed vesicles is altered during the stimulus train and is consistent with the notion that, as intracellular Ca^2+^ accumulates during repetitive stimulation, Ca^2+^-binding to the Munc13-1 C_2_B domain favors more slanted orientations of Munc13-1 (Fig. 1E) that mediate release more efficiently than the perpendicular orientations. Thus, based on this model, K603 contributes to binding of the C_1_C_2_B region to the plasma membrane in the orientations present in the absence of Ca^2+^ that mediate initial priming, but not in the slanted orientations favored upon Ca^2+^ binding to the C_2_B domain that are increasingly populated and lead to enhanced fusogenicity of the primed vesicles during the stimulus train (Fig. 1B,C, Figure 1-Figure supplement 2). In contrast R769 participates in membrane binding in both orientations, consistent with the stronger depression caused by the R769E mutation compared to the K603E mutation.

The K720E mutant displayed stronger depression than WT, based on normalized EPSC plots (Fig. 4F), but plots of absolute EPSC amplitudes show that the K720E mutant starts with higher amplitudes that depress faster over time, reaching the same steady state as WT (Fig. 4E). Thus, the stronger depression observed initially for the K720E mutant is most likely due to the initially higher release probability exhibited by this mutant (Fig. 2C) but, based on the two-state model of Fig. 1D,E, the K720E mutation does not affect release later during the stimulus training because K720 does not participate in binding in the slanted orientations favored by Ca^2+^ accumulation. In contrast, the K706E mutation led to depression that was similar to that observed for WT Munc13-1 in normalized EPSC plots (Fig. 4I), and absolute EPSC amplitudes show that the EPSCs observed for the K706E mutation were smaller than those of WT throughout the stimulus train (Fig. 4H). Since the Pvr of the K706E mutant was lower than that of WT (Fig. 2C), these observations suggest that the release probability remained low for the K706E mutant throughout the stimulus train. The effects observed for the K706E mutation are opposite to those caused by a mutation in the corresponding lysine residue of Munc13-2 to tryptophan (Shin et al., 2010) and may arise because the mutation hinders the transition to the slanted orientation, but they may also reflect a decreased fusogenicity of primed vesicles. Finally, the effects observed for the K603E/K720E and (K706E/R769E) double mutants (Fig. 5A-F) and the K603E/K720E/K706E/R769E quadruple mutants (Fig. 5G-I) appeared to be a combination of the effects caused by the single mutations, with some dominance by the K603E and the R769E mutations.

**Figure 5.**
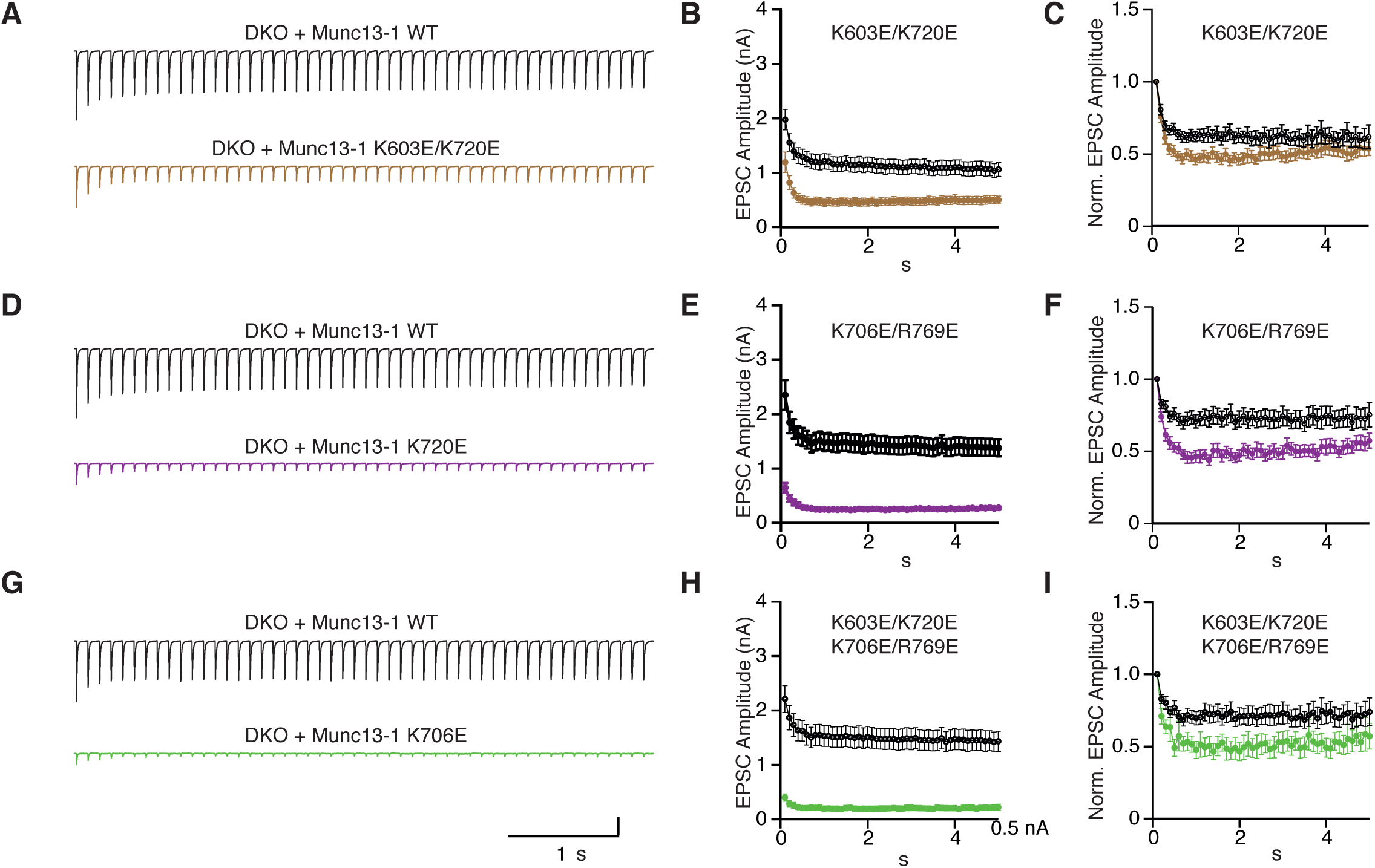
Short term plasticity behavior of combined point mutations within the Munc13-1 C_1_-C_2_B region. (**A**) Average EPSC traces recorded during a 10 Hz train of 50 action potentials and summary graphs of absolute (**B**) and normalized (**C**) EPSC amplitudes from autaptic hippocampal glutamatergic Munc13-1/2 DKO neurons expressing Munc13-1 WT (black) n=71/5 and Munc13-1 K603E/K720E (brown) n=61/5 (WT). Average EPSC traces recorded during a 10 Hz train (**D**) and summary graphs of absolute (**E**) and normalized (**F**) EPSC amplitudes from autaptic hippocampal glutamatergic Munc13-1/2 DKO neurons expressing Munc13-1 WT (black) n=77/6 and Munc13-1 K706E/R769E (purple) n=48/6 (WT). Average 10 Hz train (**G**) and summary graphs of absolute (**H**) and normalized (**I**) EPSC amplitudes from autaptic hippocampal glutamatergic Munc13-1/2 DKO neurons expressing Munc13-1 WT (black) n=65/5 and Munc13-1 K603E/K720E/K706E/R769E (green) n=17/5. Action potential artifacts were blanked.

At synapses, phorbol esters increase the release probability at least in part by activating Munc13-1 through its C_1_ domain (Basu et al., 2007; Rhee et al., 2002), mimicking the effects of DAG. The two state model predicts that this potentiation arises because DAG/phorbol esters favor the slanted orientations of Munc13-1 that facilitate SNARE complex formation and fusion. To test this notion, we analyzed the effects of the mutations in basic residues of the Munc13-1 C_1_C_2_B region on potentiation by a phorbol ester [phorbol 13,14-dibutyrate (PDBu)]. PDBu caused a robust potentiation of EPSCs in DKO autaptic cultures rescued with WT Munc13-1 (Fig. 6A,B), as expected, and the K603E and R769E mutants exhibited even higher potentiation (Fig. 6A). These results suggest that PDBu partially compensates for the priming defects caused by the K603E and R769E mutations. In contrast, the K720E mutant displayed less potentiation than WT Munc13-1, which, based on the two state model, may arise because the K720E mutation already facilitates the transition from the perpendicular to the slanted orientation and thus has an intrinsically higher release probability. The potentiation observed for the K706E mutant was similar to that observed for WT Munc13-1, suggesting that this mutation does not affect the transition between orientations but rather downstream events that lead to synaptic vesicle fusion. The K603E/K720E and K706E/R769E double mutants exhibited similar enhancements of PDBu-induced potentiation as the K603E and R769E single mutants, respectively, showing again that these mutations dominate the phenotypes of the double mutants. The results obtained for the rescue with the K603E/K720E/K706E/R769 quadruple mutant need to be interpreted with caution because this mutant exhibited a high-number of non-responsive neurons and that the EPSCs for the responsive neurons were very weak. However, it is noteworthy that we still were able to observe PDBu potentiation for this mutant.

**Figure 6.**
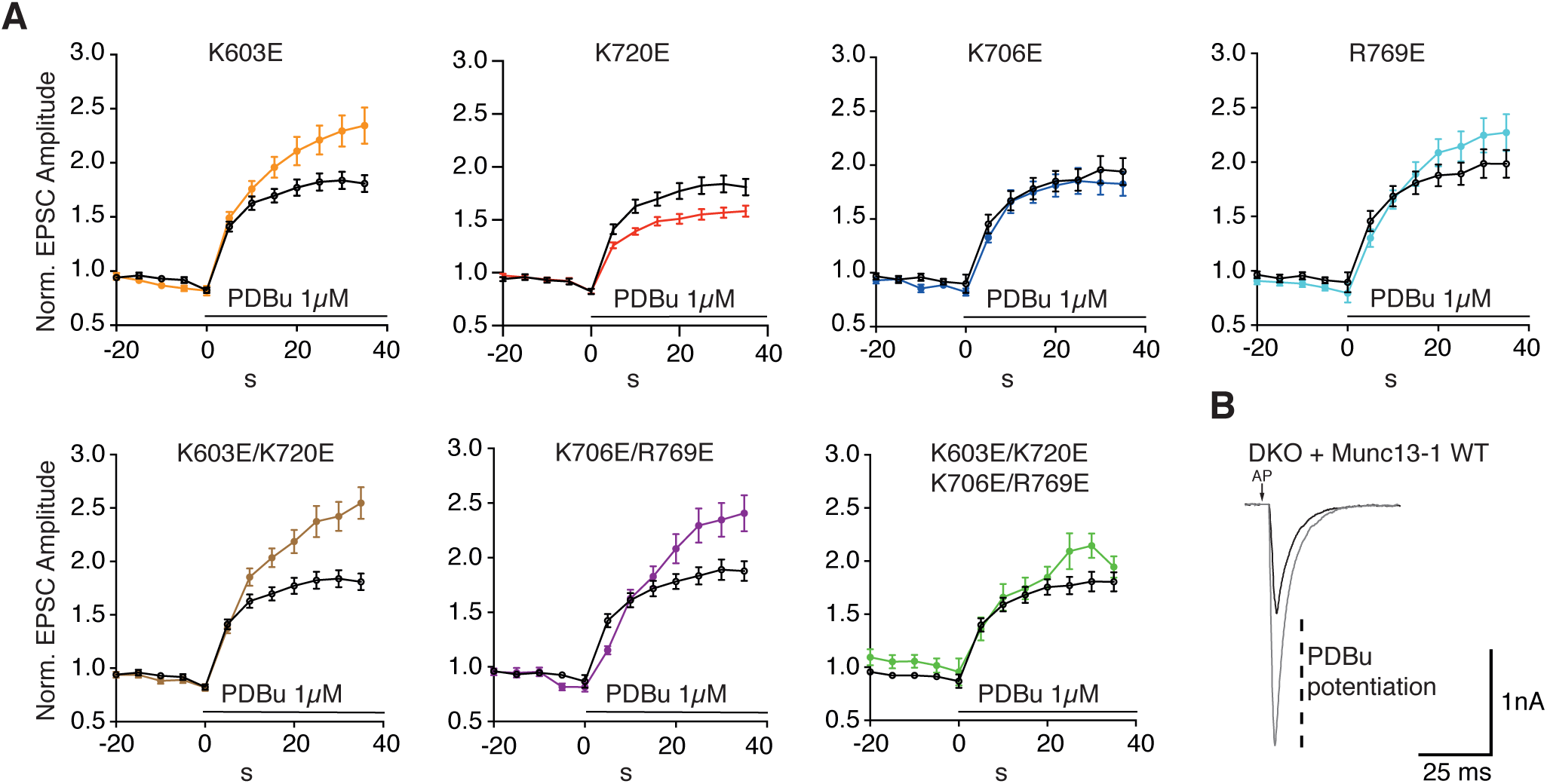
Analysis of neurotransmitter release potentiation induced by phorbol ester-activation of Munc13-1 C_1_ domain. (**A**) Summary graphs of potentiation of EPSC amplitudes by 1 μM PDBu evoked at 0.2 Hz. EPSCs were normalized to the first EPSC amplitude recorded in extracellular solution from Munc13- 1/2 DKO neurons expressing Munc13-1 WT (black) or Munc13-1 C_1_-C_2_B mutants (color code). Munc13-1 K603E (orange) n=56/5 (WT n=58/5), Munc13-1 K720E (red) n=57/5 (WT n=58/5), Munc13-1 K706E (blue) n=40/4 (WT n=38/4), Munc13-1 R769E (light blue) n=38/4 (WT n=36/4), Munc13-1 K603E/K720E (brown) n=56/5 (WT n=58/5) Munc13-1 K706E/R769E (purple) n=42/6 (WT n=57/6) and Munc13-1 K603E/K720E/K706E/R769E (green) n=27/5 (WT n=54/5). Solid symbols in the graph represent average normalized EPSC amplitudes ± SEM at each time point. (**B**) Exemplary EPSC traces from Munc13-1/2 DKO neurons expressing Munc13-1 WT before (black) and at 30 s of PDBu application (grey).

Overall, these results show that the basic residues in the Munc13-1 C_1_-C_2_B region modulate the potentiation of synaptic responses by PDBu and, together with the data obtained with repetitive stimulation, they support the notion proposed in the two state model that two faces of the C_1_-C_2_B region are critical for Munc13-1-dependent short-term plasticity.

### Mutations in the Munc13-1 C_1_-C_2_B polybasic region impair membrane binding

The proposal that C_1_C_2_BMUNC_2_C can bridge membranes in two different orientations (Fig. 1D,E) emerged from the structural studies of the C_1_C_2_BMUN fragment that revealed the polybasic face of the C_1_-C_2_B region (Xu et al., 2017), as well as from liposome clustering assays and reconstitution experiments with liposomes containing synaptobrevin (V-liposomes) and liposomes containing syntaxin-1 and SNAP-25 (T-liposomes). Thus, fusion of these liposomes in the presence of Munc18-1, NSF and αSNAP requires Ca^2+^ and the Munc13-1 C_1_C_2_BMUNC_2_C fragment, and depends on the ability of C_1_C_2_BMUNC_2_C to bridge the liposome membranes, but the liposome clustering activity of C_1_C_2_BMUNC_2_C is comparable in the absence and presence of Ca^2+^, indicating that Ca^2+^ induces a switch to an active orientation required for fusion (Liu et al., 2016; Quade et al., 2019). Mutagenesis studies demonstrated the key importance of the C_2_C domain for the ability of C_1_C_2_BMUNC_2_C to bridge membranes and support liposome fusion (Quade et al., 2019), but the role of the C_1_C_2_B region has not been tested. To address this question and attempt to rationalize the functional effects of the basic mutations in the C_1_C_2_B region observed with electrophysiology, we used a combination of assays that measure liposome binding, clustering or fusion.

We first tested the effects of these mutations on binding to liposomes with the lipid composition that we normally used for T-liposomes, which mimics that of the plasma membrane (Ma et al., 2013), using liposome co-sedimentation assays. All mutations decreased Ca^2+^-independent liposome binding to some degree compared to wild type (WT) C_1_C_2_BMUNC_2_C (top panels of Fig. 7A,B), which may arise in part because they all decrease the overall positive electrostatic potential in the region. Among the single point mutations, K603E and R769E impaired binding most strongly, while K720E had an intermediate effect and K706E exhibited the weakest effect on binding. Ca^2+^-independent liposome binding was strongly disrupted by the two double mutations and the quadruple mutation. These results are consistent with the Ca^2+^- independent binding mode predicted for the C_1_-C_2_B region (Fig. 1B, Figure 1-Figure supplement 2A), with a natural variation in the energetic contributions of the residues of the polybasic face to binding that is manifested in the different impairment caused by the K720E mutation compared with K603E and R769E. Moreover, the strong impairment of Ca^2+^-independent liposome binding caused by the K603E and R769E mutations correlate with the disruption of priming caused by these mutations (Fig. 2A), supporting the notion that Ca^2+^-independent membrane binding of the C_1_C_2_B region is important for SV priming.

**Figure 7.**
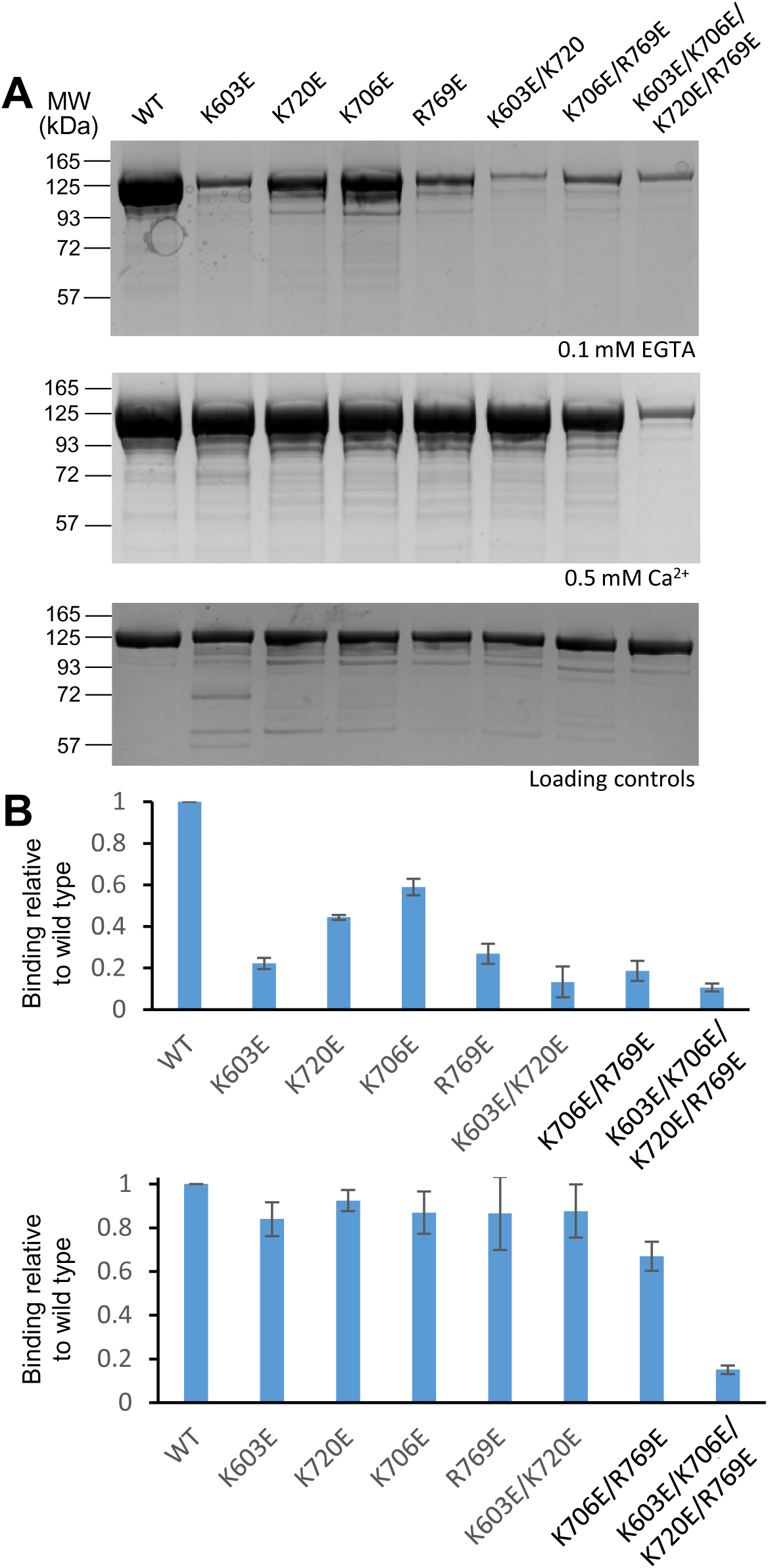
Mutations in basic residues of the C_1_-C_2_B region differentially impair the liposome affinity of the C_1_C_2_BMUNC_2_C fragment. **A.** Liposome co-sedimentation assays were performed with WT and mutant Munc13-1 C_1_C_2_BMUNC_2_C fragments and the pellets were analyzed by SDS-PAGE followed by coomassie blue staining. The top and middle images show experiments performed in the presence of 0.1 mM EGTA or 0.5 mM Ca^2+^, respectively. The bottom panel shows loading controls. The positions of molecular weight markers are indicated on the left. The total amount of C_1_C_2_BMUNC_2_C used for each sample of the co-sedimentation assays was 10 μg. Loading controls contained 2 μg of protein. **B.** Quantification of the relative amount of liposome binding of the mutant C_1_C_2_BMUNC_2_C fragments with respect to WT C_1_C_2_BMUNC_2_C. The bands of WT and mutant C_1_C_2_BMUNC_2_C fragments in gels from three independent experiments performed in the presence of 0.1 mM EGTA or 0.5 mM Ca^2+^ were quantified with ImageJ and normalize to the average value obtained for WT C_1_C_2_BMUNC_2_C. Bars indicate average values and bars show standard deviations.

Ca^2+^-dependent liposome binding was not substantially decreased by any of the single mutations or by the K603E/K720E double mutation. The K706E/R769E mutation appeared to impair Ca^2+^-dependent liposome binding to a moderate degree, while binding was almost abrogated by the quadruple mutation (Fig. 7A, middle panel, and 7B, lower panel). These results suggest that Ca^2+^ substantially increases the affinity of C_1_C_2_BMUNC_2_C for the liposomes. Such an increase may not be observable for WT C_1_C_2_BMUNC_2_C because under the conditions of our experiments binding is likely strong enough for saturation in the absence and presence of Ca^2+^, but Ca^2+^-induced enhancement of binding becomes detectable when the mutations decrease the overall affinity. For the same reason, the effects of single mutations on Ca^2+^- dependent liposome binding may not be detectable even if the mutated residues contribute to binding, but the effects observed for the double mutants support the notion that K706 and R769 contribute more to Ca^2+^-dependent binding than K603 and K720, consistent with the model of Fig. 1C, Figure 1-Figure supplement 2B. The strong effect caused by the quadruple mutation indicates that the latter residues also contribute to Ca^2+^-dependent liposome binding, likely by decreasing the overall electrostatic potential in the region.

We next analyzed the ability of the various mutants to cluster protein-free liposomes with the same lipid compositions of V- and T-liposomes using dynamic light scattering (DLS). Figure 8 compares the particle size distributions observed for a mixture of the V- and T-liposomes alone (black bars) with those observed for the same mixtures in the presence of WT or mutant C_1_C_2_BMUNC_2_C and the absence (blue bars) or presence (red bars) of Ca^2+^. The population weighted average radius calculated under each condition (Figure 8-Figure supplement 1) provides an idea of the overall amount of liposome clustering, but note that these averaged radii need to be interpreted with caution because of the difficulty in accurately calculating populations of very large particles. As observed previously (Liu et al., 2016), WT C_1_C_2_BMUNC_2_C caused strong liposome clustering, and the amount of clustering was comparable in the absence and presence of Ca^2+^ (Fig. 8A). In the absence of Ca^2+^, the liposome clustering activity was severely impaired by the single K603E and R769E mutations, the two double mutations and the quadruple mutation, whereas the single K706E and K720E mutants still supported robust liposome clustering (Fig. 8B-H). In contrast, all the mutants except the quadruple mutant induced substantial liposome clustering in the presence of Ca^2+^. These results generally correlate with the liposome binding data, and reveal that the liposome clustering activity is affected more strongly by the mutations in the absence than in the presence of Ca^2+^.

**Figure 8.**
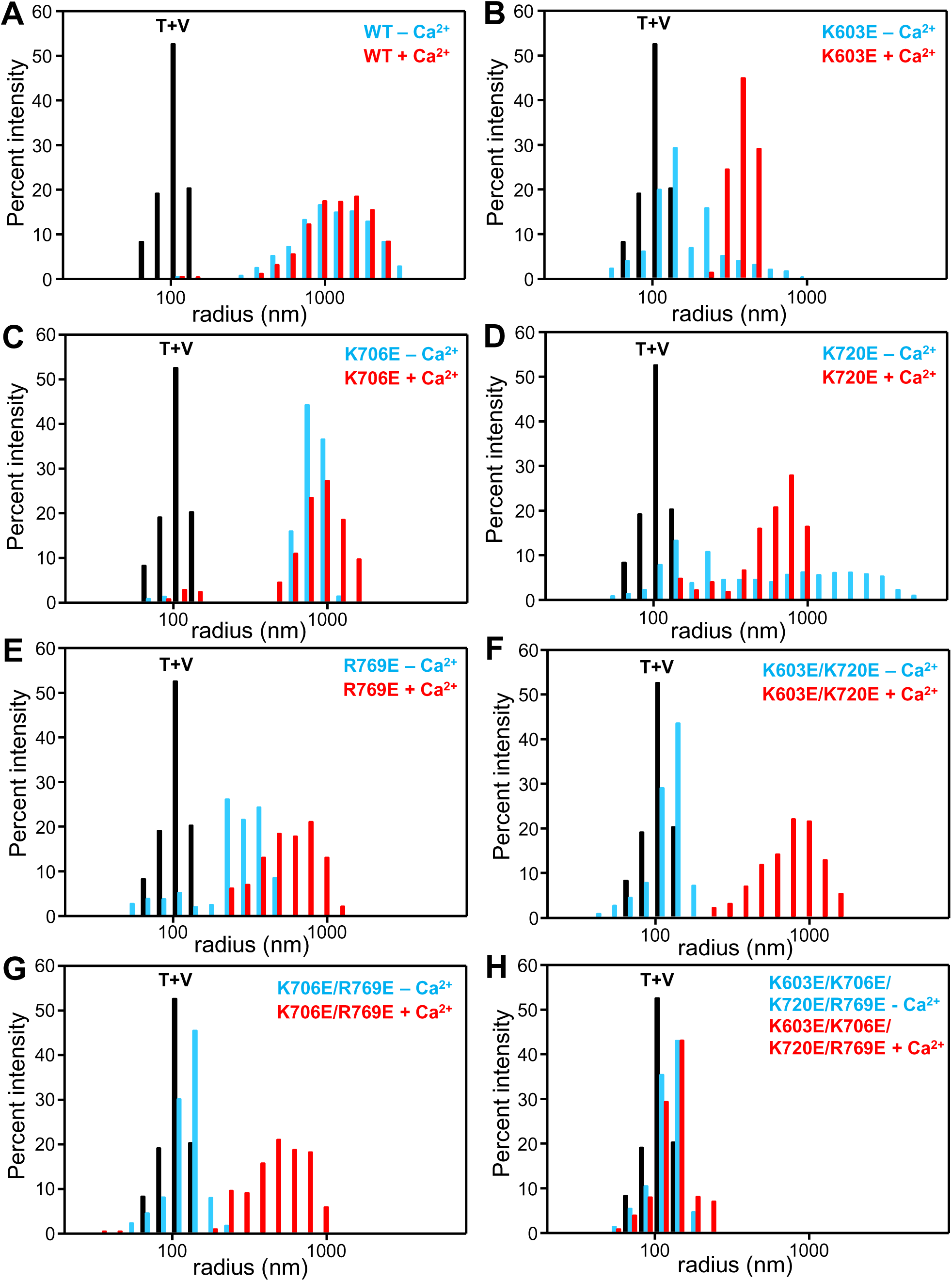
Mutations in basic residues of the C_1_-C_2_B region differentially impair the ability of the C_1_C_2_BMUNC_2_C fragment to cluster V- and T-liposomes. The ability of WT and mutant Munc13-1 C_1_C_2_BMUNC_2_C fragments to cluster V- and T-liposomes was analyzed by DLS. The diagrams show the particle size distributions observed in mixtures of the V- and T-liposomes alone (T+V, black bars), after addition of the corresponding C_1_C_2_BMUNC_2_C fragment in the presence of 0.1 mM EGTA (blue bars), and after addition of 0.6 mM Ca^2+^ to the same sample (red bars).

Overall, these data show that the C_1_-C_2_B region is critical for the liposome bridging activity of Munc13-1 C_1_C_2_BMUNC_2_C and support the notion that distinct membrane binding modes of the C_1_-C_2_B region mediate membrane bridging in the absence and presence of Ca^2+^, as predicted from the models of Fig. 1B-E. Moreover, the effects of the mutations on Ca^2+^-independent liposome clustering (Fig. 8, Figure 8-Figure supplement 1) clearly correlate with their effects on SV priming (Fig. 2A, 3A). It is noteworthy that, in the absence of Ca^2+^, the K720E mutation moderately impaired liposome binding (Fig. 7) but still allowed liposome clustering. These observations suggest that the moderate impairment of binding caused by this mutation is not sufficient to impair SV priming (Fig. 2A) but may facilitate the switch to the slanted orientation, thus enhancing evoked release (Fig. 2B) (see discussion).

### Mutations in the Munc13-1 C_1_-C_2_B polybasic region impair liposome fusion

We next turned to our reconstitution assays in which fusion between V- and T-liposomes in the presence of Munc18-1, Munc13-1 C_1_C_2_BMUNC_2_C, NSF and αSNAP is assessed by simultaneously monitoring lipid and content mixing (Liu et al., 2016). The strict Ca^2+^ dependence of fusion in these assays arises because of Ca^2+^ binding to the Munc13-1 C_2_B domain. The two state model postulates that such binding favors a more slanted orientation of C_1_C_2_BMUNC_2_C, accelerating trans-SNARE complex formation and fusion, and that this phenomenon underlies at least in part the facilitation of neurotransmitter release due to accumulation of intracellular Ca^2+^ during repetitive stimulation (Shin et al., 2010). To test whether the Ca^2+^ concentration required to activate the Munc13-1 C_2_B domain in the liposome fusion assays is consistent with this proposal, we monitored lipid mixing between V- and T-liposomes in the presence of WT Munc18-1, Munc13-1 C_1_C_2_BMUNC_2_C, NSF and αSNAP as a function of Ca^2+^ concentration. The fluorescence of the Ca^2+^-sensing dye Fluo-4 was used to measure the Ca^2+^ concentration in each point of the titration, and lipid mixing between the liposomes was monitored simultaneously from the de-quenching of Did-lipids present in the V-liposomes. We observed a steep dependence of lipid mixing on the Ca^2+^ concentration (Fig. 9A), and fitting plots of the lipid mixing observed at 1500 or 700 s as a function of Ca^2+^ concentration to a Hill equation yielded EC50s of 584 nM and 966 nM, respectively (Fig. 9B). These data show that the Munc13-1 C_2_B domain is activated in these fusion assays at submicromolar Ca^2+^ concentrations, comparable to those expected to accumulate near Ca^2+^ channels in a presynaptic active zone during repetitive stimulation.

**Figure 9.**
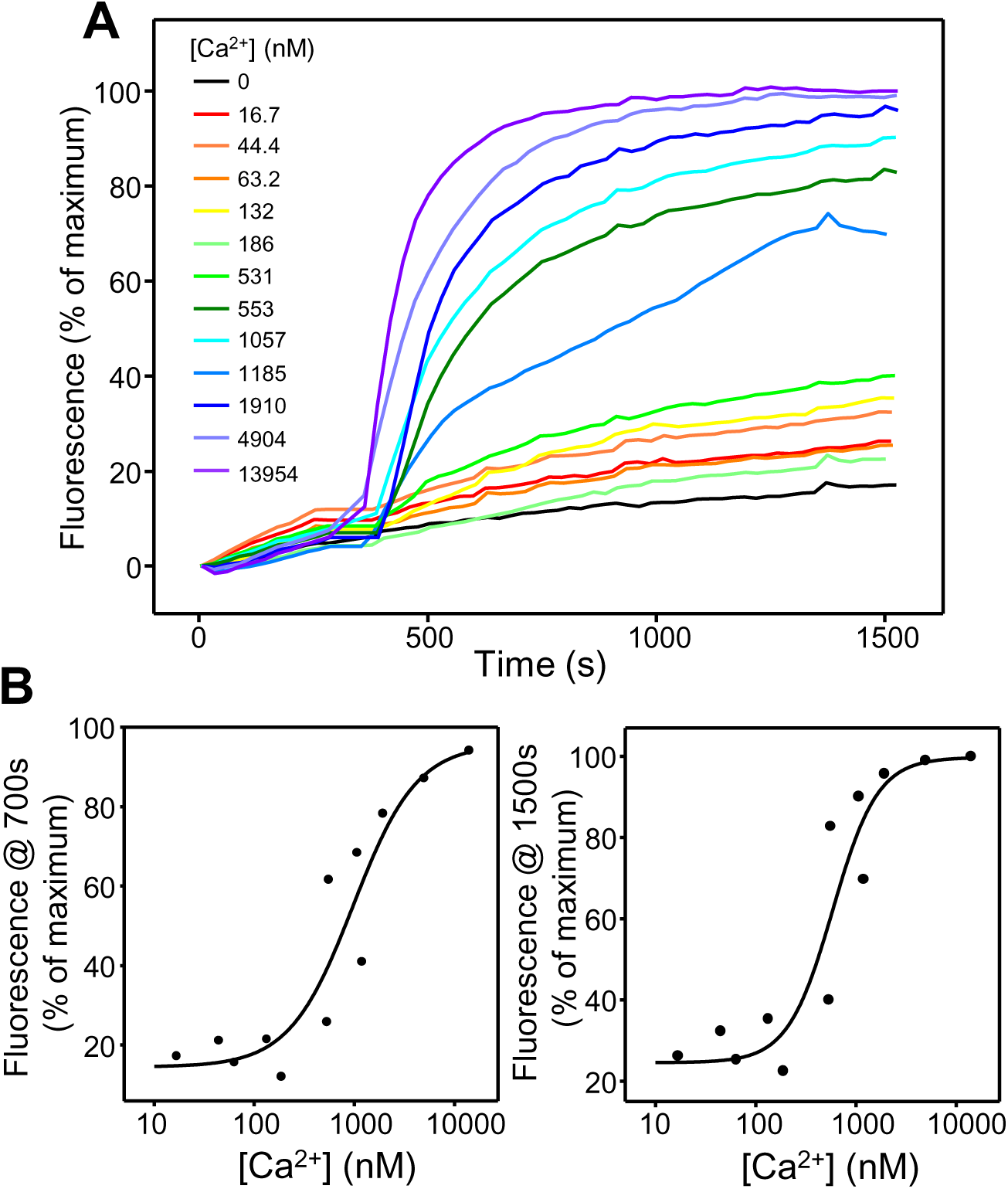
Ca^2+^-dependence of the stimulation of liposome fusion by the Munc13-1 C_1_C_2_BMUNC_2_C. **A.** Lipid mixing assays performed at different Ca^2+^ concentrations. Lipid mixing between V- and T-liposomes was measured from the fluorescence de-quenching of DiD lipids in the V-liposomes. The assays were performed in the presence of Munc18-1, NSF, αSNAP, WT Munc13-1 C_1_C_2_BMUNC_2_C fragment and Fluo-4. Experiments were started in the presence of 100 μM EGTA and Ca^2+^ was added at different concentrations after 300 s. The concentration of free Ca^2+^ in each experiment was assessed from the fluorescence of Fluo-4.

We then examined the effects of the mutations in basic residues of the C_1_-C_2_B region of Munc13-1 on liposome fusion using these assays, in which the V- and T-liposomes commonly have synaptobrevin-to-lipid ratio 1:500 and syntaxin-1-to-lipid ratio 1:800 (Liu et al., 2016). Ca^2+^-dependent liposome fusion was not impaired by any of the single point mutations or by the double K603E/K720E mutation (Fig. 10A,B,E). However, the double K706E/R769E mutation did impair fusion strongly, and fusion was abrogated by the quadruple K603E/K720E/K706E/R769E mutation (Fig. 10A,B,E). Thus, K706 and R769, which are in the Ca^2+^-binding loops of the C_2_B domain, are critical for Ca^2+^-dependent activation of fusion, while K603 and K720 did not have detectable inhibitory effects. However, it is important to note that two aspects of these assays limit their application to analyze the effects of mutations on fusion. First, inhibitory effects of the mutations on Ca^2+^-independent fusion cannot be analyzed because there is no fusion in the absence of Ca^2+^ in these assays. Second, because fusion with WT C_1_C_2_BMUNC_2_C is so efficient upon addition of Ca^2+^, gain-of-function effects cannot be observed, and moderate inhibitory effects may also be masked.

**Figure 10.**
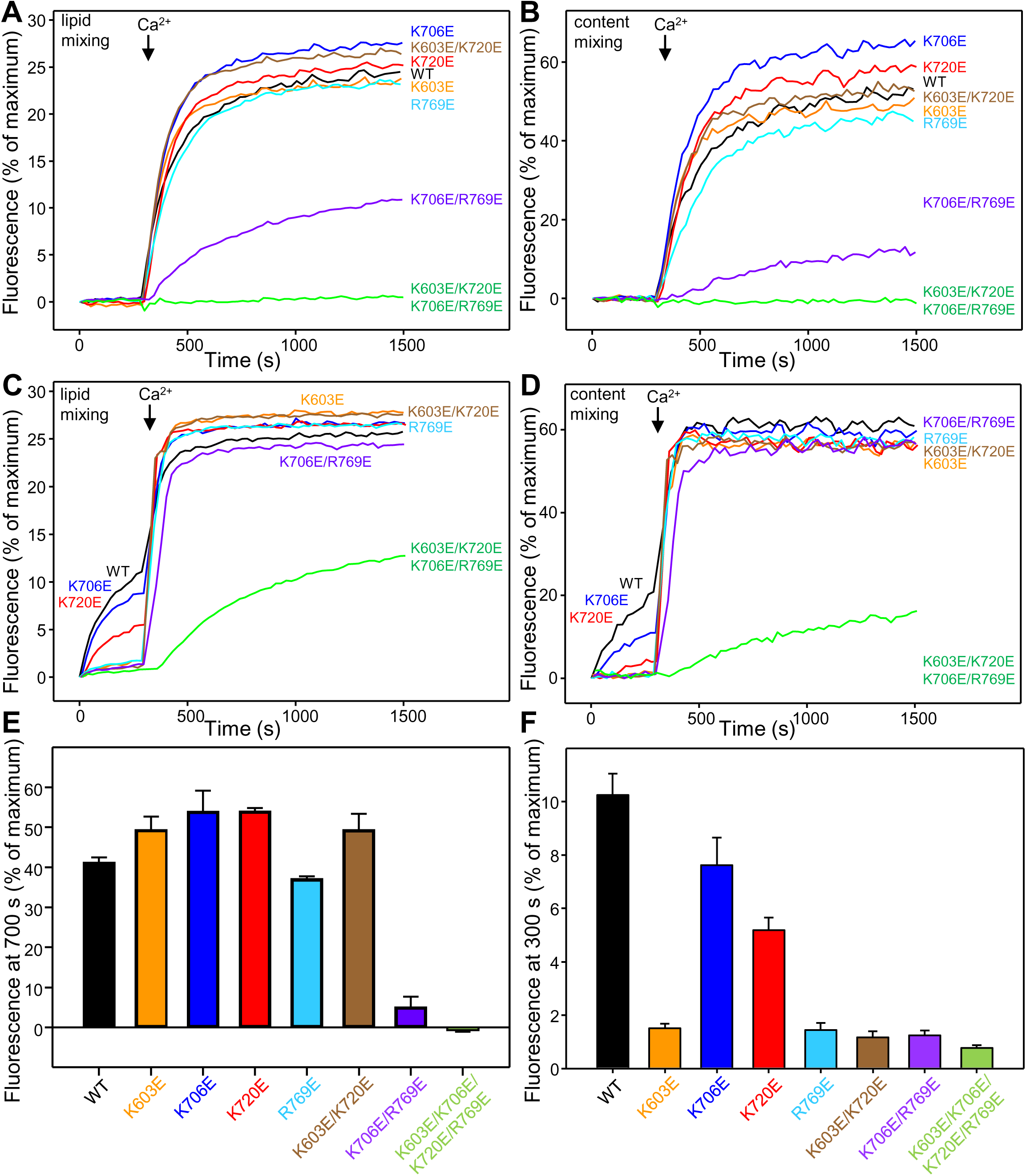
Mutations in basic residues of the C_1_-C_2_B region differentially impair the ability of the Munc13-1 C_1_C_2_BMUNC_2_C fragment to stimulate liposome fusion. **A,B.** Lipid mixing (left) between V- and T-liposomes was measured from the fluorescence de-quenching of Marina Blue-labeled lipids and content mixing (right) was monitored from the development of FRET between PhycoE-Biotin trapped in the T-liposomes and Cy5-Streptavidin trapped in the V-liposomes. The assays were performed in the presence of WT Munc18-1, NSF, αSNAP and the indicated Munc13-1 C_1_C_2_BMUNC_2_C fragments. Experiments were started in the presence of 100 μM EGTA and 5 mM streptavidin, and Ca^2+^ (600 μM) was added after 300 s. **C,D.** Analogous lipid and content mixing assays performed under the same conditions but using D326K mutant Munc18-1. **E.** Quantification of the content mixing observed after 700 s in reconstitution assays performed under the conditions of Panels **A,B**. **F.** Quantification of the lipid mixing observed after 300 s in reconstitution assays performed under the conditions of Panels **C,D**. In **E,F**, bars represent averages of the normalized fluorescence observed in experiments performed at least in triplicate, and error bars represent standard deviations.

To analyze the effects of the mutations on Ca^2+^-independent fusion, we performed analogous assays but using Munc18-1 bearing a gain-of-function mutation (D326K) that supports some Ca^2+^- independent liposome fusion (Sitarska et al., 2017) (Fig. 10C,D). The Ca^2+^-independent component of fusion, as assessed from the extent of content mixing, was somewhat inhibited by the K706E mutation, was impaired more strongly by the K720E mutation, and was abolished by all other mutations (Fig. 10D). Since lipid mixing was disrupted to a smaller extent by the mutations and hence yielded a higher dynamic range for quantification, we measured the lipid mixing at 300 s, right before Ca^2+^ addition, in repeat experiments performed under the same conditions (Fig. 10F). This analysis showed that lipid mixing was strongly impaired by all the mutations except the K706E and K720E mutations, which inhibited lipid mixing moderately. These results correlate with those obtained in the analyses of SV priming (Fig. 2A, 3A) and Ca^2+^-independent liposome clustering (Fig. 8), confirming the importance of the membrane bridging activity of C_1_C_2_BMUNC_2_C for its ability to mediate liposome fusion and supporting the notion that this activity is critical for SV priming.

To address the second issue and assess whether inhibitory or stimulatory effects of the mutations on Ca^2+^-dependent fusion in the experiments of Fig. 10A,B might not have been observed because of its high efficiency with WT C_1_C_2_BMUNC_2_C, we performed additional assays under conditions that we recently developed to render Ca^2+^-dependent fusion less efficient and highly sensitive to the Ca^2+^ sensor synaptotagmin-1 (Syt1) (Stepien and Rizo, 2021). In these assays, we monitored fusion between liposomes containing synaptobrevin and Syt1 at 1:10,000 and 1:1,000 protein-to-lipid (P/L) ratios, respectively, with liposomes containing syntaxin-1 and membrane-anchored SNAP-25 at 1:5,000 and 1:800 P/L ratios, respectively. The results that we obtained were similar to those observed under our standard conditions (Fig. 10A,B,E), as Ca^2+^-dependent fusion was markedly disrupted by the double K706E/R769E mutation, was abrogated by the quadruple K603E/K720E/K706E/R769E mutation, and was not substantially affected by the other mutations (Figure 10-Figure supplement 1). We note that the K720E mutation, which leads to enhanced neurotransmitter release (Fig. 2B), appeared to yield the most efficient Ca^2+^-dependent fusion (Figure 10-Figure supplement 1C,D) but we cannot draw firm conclusions in this respect given the variability that we normally observe in these assays (Stepien and Rizo, 2021).

Overall, the results obtained in the liposome fusion assays strongly support the proposal that binding of the C_1_-C_2_B region of C_1_C_2_BMUNC_2_C to the T-liposomes is crucial for liposome clustering and fusion, and that different binding modes involving this region in the absence and presence of Ca^2+^ underlie the drastic activation of liposome fusion observed in our assays. Moreover, our data suggest that these assays recapitulate at least to some extent the effects of the mutations on neurotransmitter release in neurons, as the K603E and R769E mutations that disrupt synaptic vesicle priming (Fig. 2A) exhibit the strongest disruption of Ca^2+^-independent fusion among the single mutations (Fig. 10B,C,F), and the impairment of Ca^2+^-dependent fusion by the double K706E/R769E mutation but not the double K603E/K720E mutation (Fig. 10A,B,E) correlates with the effects of these mutations on the vesicular release probability (Fig. 3C).

## Discussion

Research for over three decades has led to major advances in our understanding of the basic mechanisms underlying the priming and fusion of synaptic vesicles to release neurotransmitters, including the notions that Munc18-1 and Munc13-1 coordinate assembly of SNARE complexes in an NSF-αSNAP-resistant manner (Ma et al., 2013; Prinslow et al., 2019) and that this pathway enables the multiple modes of regulation of release probability during presynaptic plasticity that depend on Munc13-1 (Park et al., 2017; Sitarska et al., 2017; Stepien et al., 2019). This large multiple domain protein plays fundamental roles in this process by accelerating opening of the syntaxin-1 closed conformation (Ma et al., 2011; Yang et al., 2015) and bridging the vesicle and plasma membranes (Liu et al., 2016; Quade et al., 2019). Thus, it seems likely that Munc13-1-dependent regulation of neurotransmitter release involves at least in part alteration of these activities. Indeed, the finding that the C_1_C_2_B region has two putative faces that can bind to membranes led to an intriguing model postulating that membrane bridging by Munc13-1 in two orientations modulates neurotransmitter release (Xu et al., 2017). Here we have tested this model and examined the role of the C_1_C_2_B region in neurotransmitter release. We find that basic residues in this region play key roles in SV priming, in dictating the vesicular release probability and in modulating short-term plasticity. The differential effects of mutations in these basic residues on neurotransmitter release are mirrored in part in their effects in liposome clustering and fusion assays. Overall, these results support the notion that changes in the orientation of Munc13-1 with respect to the membranes are critical for neurotransmitter release and some forms of short-term presynaptic plasticity.

The notion that the highly conserved C-terminal region of Munc13-1 containing the C_1_, C_2_B, MUN and C_2_C domains bridges the vesicles and plasma membranes arose naturally from the discovery of its rod-like architecture, with membrane binding domains on opposite ends of the highly elongated MUN domain (Liu et al., 2016; Xu et al., 2017). At the same time, the structural studies revealed that the Munc13-1 C_1_C_2_B region contains a large polybasic face adjacent to the region containing the DAG/phorbol ester-binding site of the C_1_ domain and the Ca^2+^/PIP_2_-binding region of the C_2_B domain (Fig. 1B,C), suggesting that Munc13-1 can bind to the plasma membrane through two different faces that yield different orientations (Xu et al., 2017). This prediction is supported by our MD simulations, which indicate that membrane interactions mediated by the polybasic face involve multiple ionic interactions in an extensive surface and lead to an almost perpendicular orientation of the Munc13-1 rod with respect to the membrane. In contrast, the simulations suggest that interactions mediated by the DAG/Ca^2+^/PIP_2_- binding face involve a smaller area and are favored by Ca^2+^-phospholipid coordination and insertion of hydrophobic residues into the membrane, in addition to ionic interactions involving basic residues, leading to a very slanted orientation (Fig. 1D,E, Figure 1-Figure supplements 1 and 2). These observations support the notion that the C_1_C_2_B region binds to membranes through the larger polybasic face in the absence of Ca^2+^, adopting perpendicular orientations, whereas slanted orientations are favored by Ca^2+^ and DAG (PIP_2_ might also favor this orientation but also participates in interactions with the polybasic face).

It is important to realize that the orientation of Munc13-1 is likely dynamic, particularly in the presence of other forces. For instance, the perpendicular orientation of Munc13-1 is expected to facilitate initiation of SNARE core complex assembly but prevent full assembly, resulting in a tug-of-war between Munc13-1 and the SNAREs in the absence of Ca^2+^ that is dramatically tilted toward full SNARE complex assembly and fusion when Ca^2+^ binding to the C_2_B domain favors slanted orientations. However, the balance can also be tilted towards slanted orientations in the absence of Ca^2+^ by the energy associated with further assembly of the SNARE complex and by proteins that bind to the SNARE complex such as Syt1 and complexins (see below). Indeed, a large amount of physiological data can be explained with a model postulating that primed vesicles exist in a dynamic equilibrium between a loosely primed state (LS), in which SNARE complexes are partially assembled and Munc13-1 bridges the vesicle and plasma membrane in a perpendicular orientation, and a tightly primed state (TS) where the SNARE complex is more fully assembled and Munc13-1 bridges the membranes in a slanted orientation (Neher and Brose, 2018). In this scenario, vesicles in the TS state are much more likely to be released by an action potential than those in the LS state, and accumulation of Ca^2+^ during repetitive stimulation increases the vesicular release probability because it shifts the equilibrium toward the TS state. Note however that methods used to measure the RRP such as application of sucrose likely release vesicles in both the LS and TS states. This two state model and the proposed nature of the two membrane-binding modes of the C_1_-C_2_B region underlying the two states are strongly supported by our data.

The strong impairments of liposome binding and clustering in the absence of Ca^2+^ caused by the K603E and R769E mutations in Munc13-1 C_1_C_2_BMUNC_2_C (Fig. 7, 8) show that the polybasic face is indeed critical for Ca^2+^-independent membrane binding and membrane bridging. These mutations also caused the strongest impairments of Ca^2+^-independent liposome fusion among the single mutants (10C,D,F). The finding that the K720E mutation in the polybasic face has smaller effects on Ca^2+^-independent liposome binding, clustering and fusion suggests that K720 has a smaller energetic contribution to membrane binding than those of K603 and R769. The K706E mutation exhibited the smallest inhibitory effects on liposome binding and fusion in the absence of Ca^2+^, which correlates with the fact that K706 is not in the polybasic face.

Ca^2+^ enhanced the affinity of all the mutants for liposomes (Fig. 7) and increased the clustering activity of the mutants that had the weakest clustering ability (K603E, R769E, and the two double mutants), except the quadruple mutant (Fig. 8). Hence, although we did not observe Ca^2+^-induced increases in liposome binding and clustering for WT C_1_C_2_BMUNC_2_C because its liposome affinity and clustering activity are very high, Ca^2+^ did increase binding and clustering when these activities were impaired by mutation. These increases may arise in part because Ca^2+^ binding to the C_2_B domain enhances the overall positive electrostatic potential and in part because the Ca^2+^-dependent binding mode with slanted orientations occurs with a higher affinity than the Ca^2+^-independent binding mode involving the polybasic face. The four single mutants and the K603E/K720E double mutant supported Ca^2+^-dependent fusion with comparable efficiency as WT C_1_C_2_BMUNC_2_C (Fig. 10A,B,E) even though the Ca^2+^-dependent clustering activity of all these mutants was somewhat lower than that of WT (Fig. 8), suggesting that fusion efficiency can remain very high as long as the clustering activity is above a certain threshold. Interestingly, the double K706E/R769E mutation in the C_2_B domain Ca^2+^-binding sites disrupted Ca^2+^-dependent liposome fusion severely (Fig. 10A,B,E) even though its clustering activity is comparable to that of C1C2BMUNC2C with the K603E/K720E double mutation in the polybasic face (Fig. 8). This finding strongly supports the notion that a Ca^2+^-induced switch to a slanted orientation is critical for Ca^2+^-induced enhancement of liposome fusion.

Our electrophysiological studies support the biological relevance of at least some of the results obtained in our in vitro assays and reveal additional effects of the mutations that could not be discerned with the liposome fusion assays, providing critical evidence in support of the two state model. The observation that only the K603E and R769E mutants among the single Munc13-1 mutants exhibited severely impaired SV priming (Fig. 2A) clearly correlates with the much stronger disruption of Ca^2+^- independent liposome binding, clustering and fusion caused by these mutations, compared to the K706E and K720E single mutants (Fig. 7, 8, 10C,D,F). These results provide compelling evidence that the polybasic face of the Munc13-1 C_1_C_2_B region plays a critical function in synaptic vesicle priming. Together with the key importance of the C_2_C domain for membrane bridging and SV priming (Quade et al., 2019), these data strongly support the notion that this critical function involves bridging of the vesicle and plasma membranes by respective interactions involving the C_2_C domain and the C_1_C_2_B region of Munc13-1. The K603E and R769E mutants also exhibited similar impairments in Ca^2+^-evoked release that arose because of the defects in priming, as the Pvr remained analogous to that of WT Munc13-1 (Fig. 2B-C), and the potentiation of release by PDBu was somewhat larger for both mutants than for WT Munc13-1 (Fig. 5A). These observations suggest that binding of PDBu to the C_1_ domain can partially compensate for the defects in priming caused by these two mutations by favoring the binding mode involving the slanted orientation, which increases vesicle fusogenicity. Interestingly, we observed a considerably stronger depression during a 10 Hz stimulus train for the R769E mutant than for the R603E mutant (Fig. 4). This finding supports the notion that accumulation of Ca^2+^ during repetitive stimulation facilitates the transition to slanted orientations, which are expected to be destabilized by the R769E mutation but not by the K603E mutation.

It was surprising that the K720E mutation in the polybasic interface did not impair SV priming and enhanced Ca^2+^-evoked release as well as the Pvr, in contrast to the effects of the K603E in a nearby basic residue (Fig. 2A-C). However, as mentioned above, there is likely a delicate balance between inhibitory and stimulatory interactions within the release machinery, and some of the interactions involving the polybasic region that are important for priming initiated by a perpendicular orientation of Munc13-1 need to be released to transit to slanted orientations that support full zippering of the SNARE complex and membrane fusion. Since the K720E mutation has only a modest effect on Ca^2+^-independent liposome binding and clustering (Fig. 7, 8D), a likely explanation of our results is that the mutant retains sufficient membrane affinity to fully support priming and facilitates the transition to the more active slanted orientations, yielding a higher release probability. This model is consistent with the finding that, in contrast to the K603E mutant, the K720E mutant exhibited less potentiation of release by PDBu than WT (Fig. 6A), likely because this mutation already promotes the slanted orientations that are favored by PDBu binding to the C_1_ domain. In 10 Hz stimulus trains, the K720E mutant depressed initially more than WT Munc13-1, but later exhibited analogous EPSC amplitudes as WT (Fig. 4D-F), likely because accumulation of Ca^2+^ during repetitive stimulation already induces slanted orientations that do not involve interactions of K720 with the membrane. Nevertheless, we cannot rule out the possibility that the effects of the K720E mutation arise from a different mechanism, for instance if the mutation alters an as yet unidentified interaction of Munc13-1.

The K706E mutation did not affect SV priming, in agreement with the notion that K706 does not participate in the interactions of the polybasic interface that mediate priming, but this mutant did decrease Ca^2+^-evoked release and hence the probability of release (Fig. 2A-C). These effects are opposite to those caused by mutation of the corresponding lysine residue of Munc13-2 to Trp (the KW mutant) (Shin et al., 2010), which caused a gain-of-function, and could arise in principle because the K706E mutation hinders the transition to slanted orientations. This interpretation is consistent with the observation that EPSCs remained lower than WT for the K706E mutant during 10 Hz repetitive stimulation (Fig. 4G-I). However, the observation that PDBu potentiation was similar for this mutant and WT Munc13-1 (Fig. 6A) suggests that the effects of the K706E mutation might not be related to the transition to slanted orientations but rather to another mechanism that directly influences fusion. For instance, the Munc13-1 C_2_B domain might cause membrane perturbations analogous to those that are believed to underlie the function of the Syt1 C_2_ domains in triggering release (Fernandez-Chacon et al., 2001; Rhee et al., 2005). It is also possible that the phenotypes caused by the K706E mutation and other mutations studied here reflect effects of Munc13-1 in more than one step leading to release, which complicates the interpretation of the data.

Despite these uncertainties in the interpretation of our results, the overall data strongly support the notions that the Munc13-1 C_1_C_2_B region plays a critical role in SV priming and that this region underlies two types of interactions with the plasma membrane that yield different orientations of Munc13-1. This model is also consistent with the previous observation that a mutation that is expected to unfold the Munc13-1 C_1_ domain leads to decreased priming but increased release probability in mice (Basu et al., 2007; Rhee et al., 2002), as the absence of the C_1_ domain is expected to impair the initial binding to the plasma membrane that is important for priming but likely helps to adopt slanted orientations that facilitate release. Similarly, the finding that deletion of the C_1_ domain or the C_2_B domain of *C. elegans* unc-13 enhances release but deletion of both domains strongly impairs release (Michelassi et al., 2017) suggest that both the C_1_ domain and the C_2_B domain are important to stabilize the perpendicular orientations that mediate priming but hinder transition to the active, slanted orientations; however, at least one of the two domains needs to be present to mediate the interaction with the plasma membrane.

Clearly, further research will be necessary to better understand the nature of the LS and TS states involving different orientations of Munc13-1 and the factors that influence the equilibrium between them. For instance, the equilibrium is most likely shifted toward the TS state by Syt1, which promotes trans-SNARE complex assembly, and by complexins, which stabilize assembled trans-SNARE complexes (Prinslow et al., 2019). However, binding of Syt1 and complexins to the SNARE complex is also believed to prevent final zippering of the very C-terminus of the SNARE complex before Ca^2+^ influx, thus ensuring synchronicity of release upon Ca^2+^ influx (Voleti et al., 2020). Other proteins such as CAPS, which has functions related to those of Munc13s (Jockusch et al., 2007), may also affect the equilibrium between the loose and tight primed states. Much we have learned but much remains to be learned about this fascinating system.

## Materials and methods

### Molecular dynamics simulations

A square lipid bilayer of 19.347 nm x 19.347 nm was built in the membrane builder module in the Charm-gui (Jo et al., 2008) website (https://charmm-gui.org/). The bilayer had the following composition, which mimics that of the plasma membrane (Chan et al., 2012), except for having somewhat higher amounts of PIP_2_ and DAG (the number of molecules of each lipid is indicated; percent is indicated in parenthesis): upper leaflet, 315 (45%) cholesterol, 56 (8%) 16:0-18:1 phosphatidylcholine (POPC), 91 (13%) 18:0-22:6 phosphatidyltethanolamine, 42 (6%) 18:0-22:4 phosphatidyltethanolamine (SAPE), 70 (10%) 18:0-18:1 phosphatidylserine (SOPS), 70 (10%) 18:0-22:6 phospatidylserine (SDPS), 35 (5%) 18:0-20:4 phosphatidylinositol 4,5-bisphosphate (SAPI2D), 21 (3%) 18:0-20:4 glycerol (SAGL); lower leaflet, 315 (46.1%) cholesterol, 278 (40.6%) POPC, 42 (6.1%) SDPE, 28 (4.1%) SAPE, 21 (3.1%) SAGL. The crystal structure of the Munc13-1 C_1_C_2_BMUN fragment lacking residues 1408-1452 (which correspond to a long variable loop) (PDB code 5UE8) lacks multiple loops for which no density could be observed (Xu et al., 2017). To build a complete model of this fragment (except for residues 1408-1452), the coordinates of this crystal structure were merged with those of the missing loops in the NMR structure of the C_1_ domain (PDB code 1Y8F) (Shen et al., 2005), the crystal structure of the Ca^2+^-bound C_2_B domain (PDB code 6NYT) (Shin et al., 2010) and the refined crystal structure of the MUN domain (PDB code 5UF7) (Xu et al., 2017), after superimposing the common coordinates of these structures. Two additional missing loops were modeled with Robetta (Song et al., 2013) (https://robetta.bakerlab.org/). We placed the Ca^2+^-free C_1_C_2_BMUN model above the upper leaflet in an approximately perpendicular orientation with the polybasic face close to the membrane for the Ca^2+^-free simulation (black wire diagram in Figure 1-Figure supplement 1A). For the Ca^2+^-bound simulation, we built another system that included two Ca^2+^ ions bound to the corresponding binding sites of the C_2_B domain (Shin et al., 2010) and where C_1_C_2_BMUN was placed in a more slanted orientation such that the membrane was close to the DAG- and Ca^2+^/PIP_2_-binding sites of the C_1_ and C_2_B domains, respectively (black wire diagram in Figure 1-Figure supplement 1C).

All computations with these systems were performed with Gromacs 5.0.6 at the BioHPC supercomputing facility of UT Southwestern or with Gromacs 2019.6 at the Texas Advanced Computing Center using the CHARMM36 force field (Best et al., 2012). TIP3P explicit water boxes were built for both systems (24.3 x 26.7 x 24.3 nm^3^ for the Ca^2+^-free system; 24.3 x 28.3 x 24.3 nm^3^ for the Ca^2+^-bound system), and potassium and chloride ions were added to yield a KL concentration of 145 mM and overall charge neutrality, resulting in systems of 1.55 (Ca^2+^ free) and 1.64 (Ca^2+^-bound) million atoms. The systems were energy minimized, heated to 310 K over the course of 1 ns in the NVT ensemble and equilibrated for 1 ns in the NPT ensemble. NPT production simulations were ran for 100 ns (Ca^2+^-free system) or 86 ns (Ca^2+^- bound system) using 2 fs steps and a 1.1 nm cutoff for non-bonding interactions, and periodic boundary conditions were imposed with Particle Mesh Ewald (PME) (Darden et al., 1993) summation for long-range electrostatics. Pymol (Schrödinger, LLC) was used for molecular graphics.

### *Munc13-1/2* double knock-out

*Munc13-1/2* DKO embryos (Varoqueaux et al., 2002) on an FVB/N background were used for the generation of hippocampal neuronal cultures. *Munc13-1/2* DKO embryos were obtained by caesarean section of pregnant females from timed mating of *Munc13-1***^+/-^**/*Munc13-2***^-/-^** mice. Animal Welfare Committee of Charité – Universitätsmedizin Berlin and the Berlin state government agency for Health and Social Services approved all protocols for animal maintenance and experiments (license no. G106/20). Gender of animals used for the neuronal cultures were not distinguished.

### Hippocampal neuronal cultures

Single *Munc13-1/2* DKO hippocampal neurons seeded onto micro-islands of astrocytes (autaptic hippocampal neurons) were prepared as previously reported (Bekkers and Stevens, 1991). Briefly, HCl-cleaned 30 mm diameter glass coverslips were coated with agarose type II-A (Sigma-Aldrich) and dried for more that 48 hrs. Pre-coated coverslips were printed by stamping (0.2 mm spot diameter and 0.5 mm spot interspace) with a cell-attachment mixture of rat tail collagen (BD Biosciences), poly-D-lysine (Sigma-Aldrich) and acetic acid (XX). Astrocytes were prepared from cerebral cortices of C57BL/6N mice of either sex at postnatal day 0-1 (P0-1). Cortices were isolated and digested with 0.25% trypsin-EDTA solution (Gibco) for 15 min at 37 °C. Enzymatic solution was discarded, and cortices were washed twice with Dulbecco’s Modified Eagle Medium (DMEM) (Gibco) supplemented with 10% fetal bovine serum (FBS) (PAA) and 50 IU/ml penicillin and 50 µg/ml streptomycin (Gibco). After washing, tissue was homogenized in the same medium and cell suspension was cultured in 75 cm^2^ flasks containing DMEM supplemented with FBS and penicillin/streptomycin. After 15 days in vitro (DIV) non astrocytic cells were removed, and astrocytes were trypsinized and plated onto the microdot printed coverslips at a density of 5000 cell/cm^2^. Neurons were prepared from hippocampi of *Munc13-1/2* DKO embryos of either sex at embryonic day of 18.5 (E18.5). Hippocampi were isolated and digested for 45 min at 37 °C in DMEM containing 25 U/ml papain (Worthington), 1.65 mM L-cysteine (Sigma-Aldrich), 1 mM CaCl_2_ (Sigma-Aldrich), and 0.5 mM EDTA (Merck). Digestion was stopped replacing the enzymatic solution by DMEM containing 10% of FBS, 2.5 mg/ml albumin (Sigma-Aldrich) and 2.5 mg/ml trypsin inhibitor (Sigma-Aldrich). After enzymatic inactivation, hippocampi were mechanically dissociated in Neurobasal A (NBA) medium (Gibco) supplemented with 2% B27 (Gibco), 1% Glutamax (Gibco) and 50 IU/ml penicillin and 50 µg/ml streptomycin. Neurons were seeded at low density of 3000 cells/well onto coverslips with the astrocytic micro-island placed in 6 well plates filled with NBA medium supplemented with B27, Glutamax and penicillin/streptomycin. Autaptic cultures were kept in an incubator at 37 °C with 5% CO_2_ for 15 days before they were used for electrophysiological experiments.

### Munc13-1 lentiviral rescue constructs

*Munc13-1* control and point mutants were generated with a FLAG-tagged at the C-terminus. *Munc13-1* cDNA was constructed from rat *Unc13a* splice variant by PCR amplification (Camacho et al., 2017). The reverse primer harbors a 3xFLAG sequence (Sigma-Aldrich) to allow the expression analysis. The corresponding *Munc13-1-FLAG* PCR product was fused to a cleavage P2A (Kim et al., 2011) upstream of a nuclear localized GFP sequence (*NLS-GFP*) for identification of infected neurons. Munc13-1 C1 single point mutant (Munc13-1 K603E), Munc13-1 C2B single and double point mutants (Munc13-1 K706E, Munc13-1 K720E, Munc13-1 D757E, Munc13-1 R769E and Munc13-1 K720E/R769E) and Munc13-1 C1-C2B combined point mutants (Munc13-1 K603E/K706E, and Munc13-1 K603E/K706E/K720E/R769E), were generated in the *NLS-GFP/P2A/Munc13-1-FLAG* by PCR using primers which contained the mutations. All Munc13-1 rescue constructs engineered were confirmed by sequencing. The DNA cassette encoding *NLS-GFP/P2A/Munc13-1-FLAG* control or mutant variants were subsequently cloned into a FUWG lentiviral shuttle vector under the control of the human *synapsin-1*. Lentiviral particles were produced as described previously (Lois et al., 2002) with slight modifications. Briefly, HEK293T cells were co-transfected using polyethylenimine (PEI) with FUGW shuttle vector and two packaging plasmids pCMVd8.9 and pVSV-G. Cells were incubated for 72 h at 32 °C and 5% CO_2_ in NBA medium supplemented with B27 and penicillin/streptomycin. NBA medium containing lentiviruses were harvested, passed through 0.45 μm filter, concentrated by centrifugation using an Amicon tube (Ultra-15, Ultracel-100 kDa) and stored at -80 °C. Neurons were infected 24 h after plating with 50 µl of the different concentrated lentiviral rescue constructs per 35 mm diameter well.

### Electrophysiology on autaptic hippocampal neuronal cultures

Whole-cell voltage-clamp recordings were made from excitatory hippocampal autaptic neurons at room temperature after 15-18 days in vitro. Glass coverslip containing the autaptic neuronal cultures were immersed in extracellular saline solution consisting of (in mM): 140 NaCl, 2.4 KCl, 10 HEPES, 10 glucose, 2 CaCl_2_ and 4 MgCl_2_ (300 mOsm pH, 7.4). Borosilicate patch-pipette with resistances from 3-4 MΩ were filled with a KCl-based intracellular solution containing (in mM): 136 KCl, 17.8 HEPES, 1 EGTA, 4.6 MgCl_2_, 2 mM Na_2_ATP, 0.3 Na_2_GTP, 12 creatine phosphate, and 50 U/ml phosphocreatine kinase (300 mOsm, pH 7.4). Only single autaptic neurons with a nuclear localized GFP signal were used for recording. Neurons were voltage clamped at -70 mV using a Multiclamp 700B amplifier and signals were digitized using a Digidata 1440A digitizer (Molecular Devices). Series resistance was compensated at 70 %. Only neurons with access resistance of < 10 MΩ were used. Glutamatergic neurons were identified by their characteristic EPSC time course and by sensitivity of EPSC to application of glutamate receptor antagonist kynurenic acid (Tocris). Synaptic currents were acquired at 10 kHz using pClamp 10.2 software (Molecular Devices) and filtered at 3 kHz. Miniature excitatory postsynaptic currents (mEPSCs) were recorded for periods of at least 30 s in extracellular solution. Recordings in 3 mM Kynurenic acid were used to subtract false-positive events. Miniature events were detected using a template-based mEPSC algorithm implemented in Axograph X (AxoGraph Scientific). Excitatory postsynaptic currents (EPSCs) were induced by a 2 ms somatic depolarizing from -70 to 0 mV at a frequency of 0.2 Hz. The ready releasable pool (RRP) was determined by integrating the transient postsynaptic current induced by a 5 s application of 0.5 M hypertonic sucrose solution. *Pvr* was estimated as the ratio of the EPSC charge to the RRP charge. Short term plasticity was evaluated by a Pair-pulse protocol which 2 consecutives APs at 40 Hz or by a train of 50 APs at 10 Hz. Paired-pulse ratios were calculated by dividing the amplitude of the second EPSC to the first. PDBu induced potentiation was calculated by comparing the EPSC amplitude after PDBu application with the preceding EPSC amplitudes in extracellular solution.

### Immunocytochemistry

*Munc13-1/2* DKO hippocampal mass culture neurons were plated at a density of 25,000 cells, infected with the Munc13-1 C_1_-C_2_B polybasic face mutants and fixed after 15 DIV with 4% paraformaldehyde. After fixation, neurons were permeabilized, blocked and incubated overnight with rabbit polyclonal antibody against Munc13-1 (1:500; Synaptic System 126103), chicken polyclonal antibody against MAP2 (1:2,000; Merck Millipore AB5543) and guinea pig polyclonal antibody against VGLUT1 (1:4,000; Synaptic System, 135304). Primary antibodies were labelled with Alexa Fluor 488 AffiniPure donkey anti-guinea pig IgG, Rhodamine Red-X AffiniPure donkey anti-chicken IgG and Alexa Fluor 647 AffiniPure donkey anti-rabbit IgG (1:500; Jackson ImmunoResearch) for 1 hour at RT. Coverslips were mounted with Mowiol 4–88 antifade medium (Polysciences Europe). Fixed neurons were imaged on a confocal laser-scanning microscope Leica TCS SP8 with identical settings used for all samples. Neuronal cultures were visualized using a 63 × oil immersion objective. Images were acquired using Leica Application Suite X (LAsX) software at 1,024 × 1,024 pixels resolution using a *z*-series projection of 10-12 images with 0.3 μm depth intervals. Six independent neurons per group for each cultured and two different cultures were imaged and analyzed using ImageJ software.

### Electrophysiological Data Analysis and Statistics

Electrophysiological data were analyzed offline using Axograph X software version 1.4.3 (AxoGraph Scientific). Statistical analyses were carried out with GraphPad Prism software version 8 (GraphPad software). Number of neurons and cultures used for the statistical analyses are specified within the bar plots. Data was tested for normality by a D’Agostino-Pearson test showing a non-normal distribution. Electrophysiological data are presented as normalized means to the corresponding WT control group ± standard error for means (SEM), except for the Pvr and PPR that are reported as absolute means ± SEM. Statistical comparison between the different mutant groups was performed with Mann–Whitney *U* test. The significance level was set at *p* = 0.05.

#### Plasmids and recombinant proteins

Plasmids used to express the following proteins, as well as methods for expression and purification in bacteria were described previously: full-length *Homo sapiens* SNAP-25A with and without its four cysteines mutated to serine, full-length *Rattus norvegicus* synaptobrevin-2, full-length *Rattus norvegicus* Munc18-1 WT and D326K, full-length *Rattus* syntaxin-1A, full-length *Cricetulus griseus* NSF V155M mutant, *Rattus norvegicus* synaptotagmin-1 57-421 C74S/C75A/C77S/C79I/C82L/C277S (a kind gift from Thomas Söllner), and full-length *Bos taurus* α-SNAP in *E.* coli were described previously (Chen et al., 2006; Dulubova et al., 2007; Dulubova et al., 1999; Liu et al., 2016; Ma et al., 2013; Prinslow et al., 2019; Sitarska et al., 2017; Stepien and Rizo, 2021). A plasmid for bacterial expression of *Rattus Norvegicus* His6-Munc13-1 residues 529-1735 lacking a large variable loop (residues 1408-1452) to improve solubility (Ma et al., 2011) was described previously (Quade et al., 2019). Standard PCR-based recombinant DNA techniques with custom-designed primers based on the parent DNA were used to create expression vectors for the following mutants of Munc13-1 C_1_C_2_BMUNC_2_C: K603E, K706E, K720E, R769E, K603E/K720E, K706E/R769E, K603E/K706E/K720E/R769E.

Expression of His6-Munc13-1 C_1_C_2_BMUNC_2_C (WT and mutants) encoded in a pET28a vector was performed in *E. coli* BL21 (DE3) cells. Transformed cells were grown in the presence of 50 μg/ml kanamycin to an OD600 of ∼0.8 and induced overnight at 16 °C with 500 μM IPTG. Cells were harvested by centrifugation and re-suspended in 50 mM Tris, pH 8, 250 mM NaCl, 1 mM TCEP, 10% glycerol (v/v) prior to lysis. Cell lysates were centrifuged for 30 minutes at 48,000 x g to clarify the lysate and then incubated with Ni-NTA resin for 30 minutes at room temperature. The resin was washed with re-suspension buffer and re-suspension buffer with an additional 750 mM NaCl to remove contaminants. Nuclease treatment was performed on the beads for 1 hour at room temperature using 250 U of Pierce Universal Nuclease (Thermo Scientific) per liter of cells. Protein was eluted using re-suspension buffer with 500 mM imidazole and dialyzed against 50 mM Tris, pH 8, 250 mM NaCl, 1 mM TCEP, 2.5 mM CaCl, 10% glycerol (v/v), overnight at 4°C in the presence of thrombin. The solution was re-applied to Ni-NTA resin to remove any uncleaved protein and diluted twentyfold with 20 mM Tris, pH 8, 1 mM TCEP, 10% glycerol (v/v). Diluted protein was subjected to anion exchange chromatography using a HiTrapQ HP column (GE Life Sciences) and eluted in 20 mM Tris, pH 8, 1 mM TCEP, 10% glycerol (v/v) with a linear gradient from 1% to 50% of 1 M NaCl. Fractions containing protein were applied to a Superdex 200 column using 20 mM Tris, pH 8, 250 mM NaCl, 1 mM TCEP, 10% glycerol (v/v).

#### Liposome clustering assays

To prepare phospholipid vesicles, 1-palmitoyl-2-oleoyl-sn-glycero-3-phosphocholine (POPC), 1,2- dioleoyl-sn-glycero-3-phospho-L-serine (DOPS), 1-palmitoyl-2-oleoyl-sn-glycero-3-phosphoethanolamine (POPE), L-a-Phosphatidylinositol-4,5-bisphosphate (PIP2), 1-palmitoyl-2-oleoyl-*sn*-glycerol (DAG), and cholesterol dissolved in chloroform were mixed at the desired ratios and then dried under a stream of nitrogen gas. T-type liposomes contained 38% POPC, 18% DOPS, 20% POPE, 2% PIP2, 2% DAG, and 20% Cholesterol, and V-type liposomes contained 39% POPC, 19% DOPS, 22% POPE, and 20% Cholesterol. The dried lipids were left overnight in a vacuum chamber to remove the organic solvent. The next day the lipid films were hydrated with 25 mM HEPES, pH 7.4, 150 mM KCl, 10% glycerol (v/v), and vortexed for 5 minutes followed by five freeze-thaw cycles. Large unilamellar vesicles were prepared by extruding the hydrated lipid solution through a 100 nm polycarbonate filter 31 times with an Avanti Mini-Extruder. Liposome clustering induced by Munc13 fragments was analyzed using a Wyatt Dynapro Nanostar dynamic light scattering instrument (Wyatt Technology) equipped with a temperature controlled microsampler as previously described (Liu et al., 2016). Particle size was measured with T-liposomes (250 μM total lipid) and V-liposomes (125 μM total lipid) diluted in 25 mM HEPES, pH 7.4, 150 mM KCl, 10% glycerol (v/v) with 100 μM EGTA, two minutes after the specified Munc13-1 fragment (500 nM) was added to the mixture, and three minutes after the addition of 600 μM Ca^2+^. All incubations were performed at room temperature.

#### Liposome co-sedimentation assays

Liposome co-sedimentation assays were performed as described with some modifications (Quade et al., 2019; Shin et al., 2010). Briefly, lipid mixtures containing 38% POPC, 18% DOPS, 19% POPE, 2% PIP2, 2% DAG, 20% cholesterol, and 1% Rhodamine-PE were dried under a stream of nitrogen gas and kept under vacuum overnight. The next day the lipid film was re-suspended in buffer (25 mM Hepes, pH 7.4, 150 mM KCl, 1 mM TCEP, 500 mM sucrose), frozen and thawed five times, and then extruded through a 100 nm polycarbonate filter 31 times. Liposomes were diluted in sucrose-free buffer and spun at 160,000 x g for 30 min to pellet heavy liposomes. The supernatant was removed and liposomes were re-suspended in sucrose-free buffer. Liposomes were then pelleted at 17,000 x g and re-suspended in sucrose free buffer two more times. The final liposome concentration was estimated based on the absorbance of Rhodamine-PE in a known liposome sample. Liposome solutions containing 2 mM lipids and 2 μM protein were incubated for 30 min at room temperature. The liposomes and bound protein were pelleted by centrifugation at 17,000 x g for 20 min. The supernatant was removed and the liposomes were re-suspended in buffer. Re-suspended samples were boiled for 5 min and analyzed by SDS-PAGE and coomassie blue staining.

#### Liposome fusion assays

Liposome lipid and content mixing assays were performed as previously described (Liu et al., 2016; Liu et al., 2017; Stepien and Rizo, 2021). To prepare the phospholipid vesicles, POPC, DOPS, POPE, PIP2, DAG, 1,2-dipalmitoyl-*sn*-glycero-3-phosphoethanolamine-N-(7-nitro-2-1,3-benzoxadiazol-4-yl) (ammonium salt) (NBD-PE), 1,2-Dihexadecanoyl-*sn-*glycero-3-phosphoethanolamine (Marina Blue DHPE), and cholesterol in chloroform were mixed at the desired ratio and dried under a stream of nitrogen gas. T-liposomes contained 38% POPC, 18% DOPS, 20% POPE, 2% PIP2, 2% DAG, and 20% Cholesterol, and V-liposomes contained 39% POPC, 19% DOPS, 19% POPE, 20% Cholesterol, 1.5% NBD PE, and 1.5% Marina Blue DHPE. The dried lipids were left overnight in a vacuum chamber to remove the organic solvent. The next day the lipid films were hydrated with 25 mM Hepes, pH 7.4, 150 mM KCl, 1 mM TCEP, 2% n-Octyl-β-D-glucoside (β-OG) and 10% glycerol (v/v) by vortexing for 5 minutes. For the experiments of Fig. 8, 9, rehydrated lipids for T-liposomes were mixed with protein and dye to get a final concentration of 4 mM lipid, 5 μM full-length syntaxin-1, 25 μM full-length SNAP-25 with its four cysteines mutated to serine, and 4 μM R-phycoerythrin biotin-XX conjugate (Invitrogen). Rehydrated lipids for V-liposomes were mixed with protein and dye to get a final concentration of 4 mM lipid, 8 μM full-length synaptobrevin, and 8 μM Cy5-streptavidin conjugate (Seracare Life Sciences Inc.). Lipid mixtures were dialyzed 1 hr, 2 hr and overnight at 4°C in 25 mM Hepes, pH 7.4, 150 mM KCl, 1 mM TCEP, 10% glycerol (v/v) in the presence of Amberlyte XAD-2 beads (Sigma) to remove the detergent and promote the formation of proteoliposomes. The next day the proteoliposomes were harvested and mixed with Histodenz (Sigma) to a final concentration of 35%. Proteoliposome mixtures were added to a centrifuge tube with 25% Histodenz and 25 mM Hepes, pH 7.4, 150 mM KCl, 1 mM TCEP, 10% glycerol layered on top. The proteoliposomes were spun at 4°C for 1.5 hours at 55,000 RPM in an SW-60 TI rotor and the top layer was collected. Concentrations of the final T-proteoliposomes were measured by the Stewart method (Stewart, 1980). V-proteoliposome concentrations were estimated from the UV-vis absorption using a standard curve made using known quantities of liposomes containing 1.5% NBD-PE. For the experiments of Figure 10-Figure supplement 1, we used the same protocol except for the following modifications. We used WT SNAP-25 that was dodecylated as described (Stepien and Rizo, 2021) and incorporated into the T-liposomes with P:L ratio 1:800 instead of SNAP-25 with its four cysteines mutated to serine, and syntaxin-1 was incorporated into the T-liposomes with P:L ratio 1:5,000. Instead of V-liposomes, we used VSyt1-liposomes containing 40% POPC, 6.8% DOPS, 30.2% POPE, 20% Cholesterol, 1.5% NBD PE, and 1.5% Marina Blue DHPE, synaptobrevin (1:10,000 P:L ratio) and synaptotagmin-1(57-421) C74S/C75A/C77S/C79I/C82L/C277S (P:L ratio 1,1000), as described (Stepien and Rizo, 2021).

To perform the fusion assays, T-liposomes (250 μM total lipid) were first incubated with 1 μM Munc18-1 (WT or D326K as indicated), 0.8 μM NSF, 2 μM αSNAP, 2 mM ATP, 2.5 mM Mg^2+^, 5 μM streptavidin, and 100 μM EGTA for 15-25 minutes at 37°C, and then were mixed with V-liposomes or VSyt1-liposomes (125 μM total lipid), 1 μM SNAP-25 (only for the experiments of Figs. 8 and 9), and 0.1 μM WT or mutant C_1_C_2_BMUNC_2_C (except the experiments of Fig. 9C,D,F, in which the C_1_C_2_BMUNC_2_C concentration was 0.5 μM). After 5 minutes 0.6 mM Ca^2+^ was added to stimulate fusion, and 1% β-OG was added after 25 minutes to solubilize the liposomes. The fluorescence signals from Marina Blue (excitation at 370 nm, emission at 465 nm) and Cy5 (excitation at 565 nm, emission at 670 nm) were recorded to monitor lipid and content mixing, respectively. The lipid mixing data were normalized to the maximum fluorescence signal observed upon detergent addition. The content mixing data were normalized to the maximum Cy5 fluorescence observed after detergent addition in control experiments without external streptavidin.

#### Measuring Ca^2+^ concentrations during liposome fusion assays

Liposome lipid mixing assays were performed basically as described above for the experiments with V- and T-liposomes, except that we used different fluorescent lipids. To prepare the phospholipid vesicles, POPC, DOPS, POPE, PIP2, DAG, and cholesterol in chloroform and 1,1’-dioctadecyl-3,3,3’,3’-tetramethylindodicarbocyanine, 4-chlorobenzenesulfonate salt (DiD, Invitrogen) in DMSO were mixed at the desired ratio and dried under a stream of nitrogen gas. T-liposomes contained 38% POPC, 18% DOPS, 20% POPE, 2% PIP2, 2% DAG, and 20% Cholesterol, and V-liposomes contained 38.5% POPC, 19% DOPS, 19% POPE, 20% Cholesterol, and 3.5% DiD. The dried lipids were left overnight in a vacuum chamber to remove the organic solvent. The next day the lipid films were hydrated with 25 mM Hepes, pH 7.4, 150 mM KCl, 1 mM TCEP, 2% n-Octyl-β-D-glucoside (β-OG) and 10% glycerol (v/v) by vortexing for 5 minutes. Rehydrated lipids for T-liposomes were mixed with protein to get a final concentration of 4 mM lipid, 5 μM full-length syntaxin-1, 25 μM full-length SNAP-25 with the four cysteines mutated to serine. Rehydrated lipids for V-liposomes were mixed with protein to get a final concentration of 4 mM lipid and 8 μM full-length synaptobrevin. Lipid mixtures were dialyzed 1 hr, 2 hr and overnight at 4°C in 25 mM Hepes, pH 7.4, 150 mM KCl, 1 mM TCEP, 10% glycerol (v/v) in the presence of Amberlyte XAD-2 beads (Sigma) to remove the detergent and promote the formation of proteoliposomes.

To perform the fusion assays, T-liposomes (250 μM total lipid) were first incubated with 1 μM Munc18-1 wild type, 0.8 μM NSF, 2 μM αSNAP, 2 mM ATP, 2.5 mM Mg^2+^, 5 μM streptavidin, 100 uM EGTA, and 0.5 μM Fluo-4, pentapotassium salt for 15-25 minutes at 37°C. To initiate the reaction preincubated T-liposomes were mixed with V-liposomes (125 μM total lipid), 1 μM SNAP-25, and 0.1 μM C1C2BMUNC2C. After 5 minutes Ca^2+^ was added to stimulate fusion, and 1% β-OG was added after 25 minutes to solubilize the liposomes. Throughout the reaction lipid mixing was monitored using DiD de-quenching with excitation at 560 nm and emission measured at 670 nm. After solubilization of samples, the Fluo-4 emission intensity was measured from 505 to 540 nm with excitation at 465 nm. To measure the maximum fluorescence of Fluo-4 in each sample, 10 mM Ca^2+^ was added to each sample and the fluorescence spectrum was acquired again. Ca^2+^ concentrations were calculated as previously described (Grynkiewicz et al., 1985).

## Acknowledgments

We thank Berit Söhl-Kielczynski, Bettina Brokowski, Katja Pötschke, Sabine Lenz and Heike Lerch for excellent technical support, and Milo Lin for critical reading of the manuscript. We also thank the Charité Viral core facility for virus production and characterization, and Neurocure imaging core facility at the Charité Campus Mitte for services. The authors acknowledge the Texas Advanced Computing Center (TACC) at The University of Texas at Austin for providing high performance computing resources that have contributed to the research results reported within this paper (URL: http://www.tacc.utexas.edu). This research was also used computational resources provided by the BioHPC supercomputing facility located in the Lyda Hill Department of Bioinformatics, UT Southwestern Medical Center, TX (URL: https://portal.biohpc.swmed.edu). Bradley Quade was supported by NIH Training Grant T32 GM008297. This work was supported by grant I-1304 from the Welch Foundation (to JR), by NIH Research Project Award R35 NS097333 (to JR), by the German Research Council Grant CRC 958 and Reinhart Koselleck project (to CR).

## Data availability

Source data files are provided for all data figure panels.

## Supplementary Figures

**Figure 1-Figure supplement 1.**
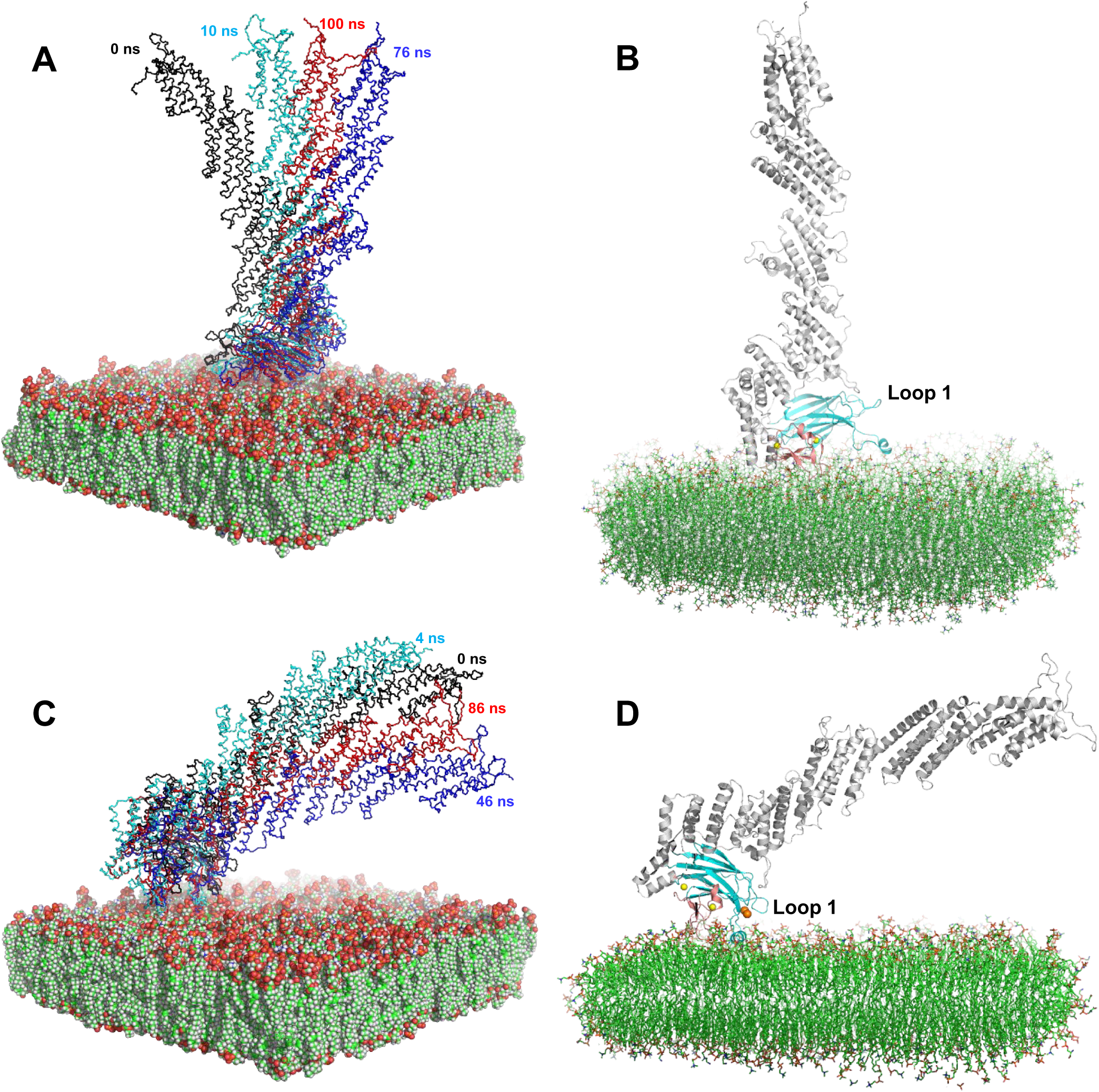
Representative conformations showing the orientations of the C_1_C_2_BMUNC_2_C fragment with respect to the flat bilayer visited during the MD simulations. **A,C**. The panels show the initial flat bilayer in space-filling models and cartoon representations of the backbone of the C_1_C_2_BMUN fragment in the initial (black) and final (red) orientation of the MD simulations performed in the absence (**A**) or presence (**C**) of Ca^2+^. **A** also shows cartoon representations of the backbone of the C_1_C_2_BMUN fragment in the two extremes of the range of orientations visited from 10 to 100 ns of the simulation. Note that the final orientation is approximately in the middle of this range. **C** also shows cartoon representations of the backbone of the C_1_C_2_BMUN in the two extremes of the range of orientations visited during the entire simulation, but we note that the orientation of C_1_C_2_BMUN was most of the time ranging between the initial orientation and the orientation observed at 46 ns. The final orientation is approximately in the middle of that range. **B,D.** Final frames of the MD simulations performed in the absence (**B**) and presence (**D**) of Ca^2+^, which are representative of the average orientations observed during the respective simulations. The C_1_C_2_BMUN fragment is shown as a ribbon diagram with the C_1_ domain in salmon, the C_2_B domain in cyan and the rest of the molecule in grey. The flay bilayer is represented by stick models. Carbon atoms are in green, oxygen atoms in red, nitrogen atoms in blue and phosphorous atoms in orange.

**Figure 1-Figure supplement 2.**
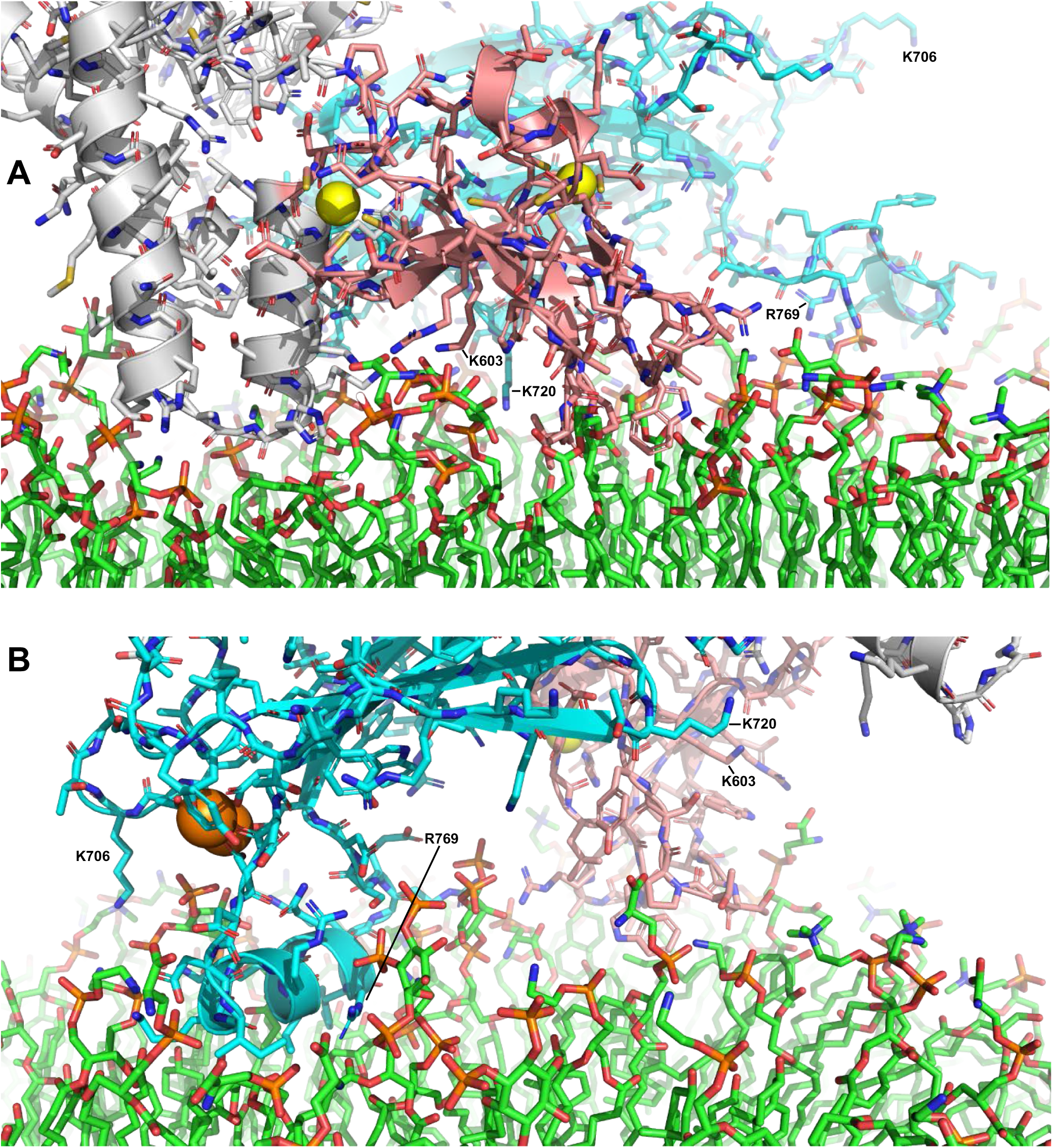
Close up views of the membrane-binding modes of the C_1_C_2_BMUN fragment observed in the final frames of the MD simulations performed in the absence (**A**) or presence (**B**) of Ca^2+^. The C_1_C_2_BMUN is shown with the backbone represented by a ribbon diagram (C_1_ domain in salmon, C_2_B domain in cyan and the rest of the backbone in grey) and the side chains represented by stick models. The bilayer is represented by stick models. Carbon atoms are in green, oxygen atoms in red, nitrogen atoms in blue and phosphorous atoms in orange. Zn^2+^ and Ca^2+^ ions are shown as yellow and orange spheres, respectively. The positions of the side chains that were mutated in this study are indicated.

**Figure 2-Figure supplement 1.**
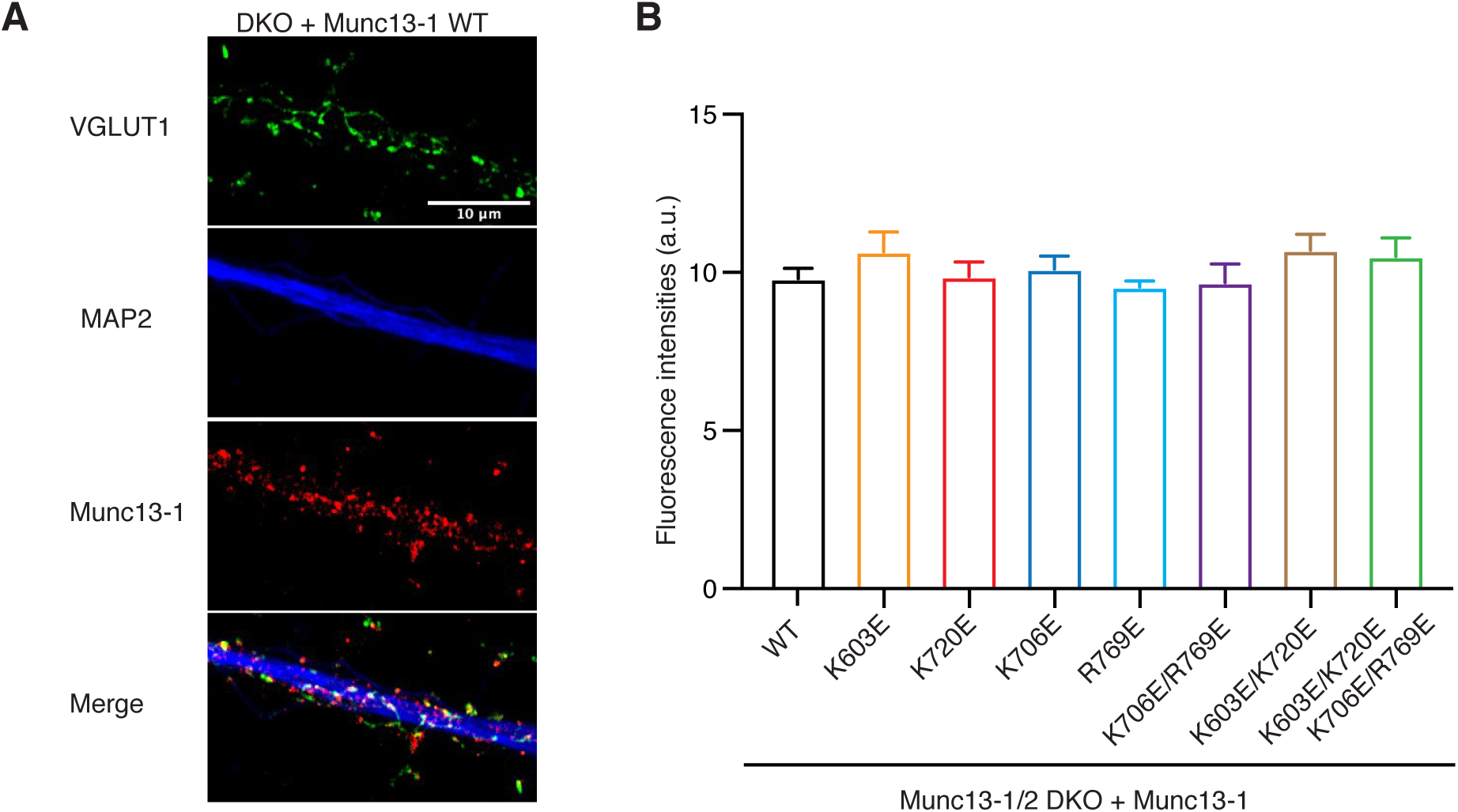
Presynaptic expression levels of Munc13-1 C_1_-C_2_B region mutants. **A.** Representative confocal images from Munc13-1/2 DKO rescued with Munc13-1 WT hippocampal neurons. Neurons were fixed and counterstained with VGLUT1, MAP2 and Munc13-1. Scale bar, 10 µm. **B.** Plot of Munc13-1 fluorescence intensity levels of DKO neurons rescued with the different mutants indicated on the x-axis. Munc13-1 mean intensity values were analyzed in 50 positives VGLUT1 puncta per cell, in 6 different cells per group and in 2 independent cultures. Significances and p-values were determined using ruskal–Wallis one-way analysis of variance followed by a multiple comparison Dunn’s *post hoc* test.

**Figure 2-Figure supplement 2.**
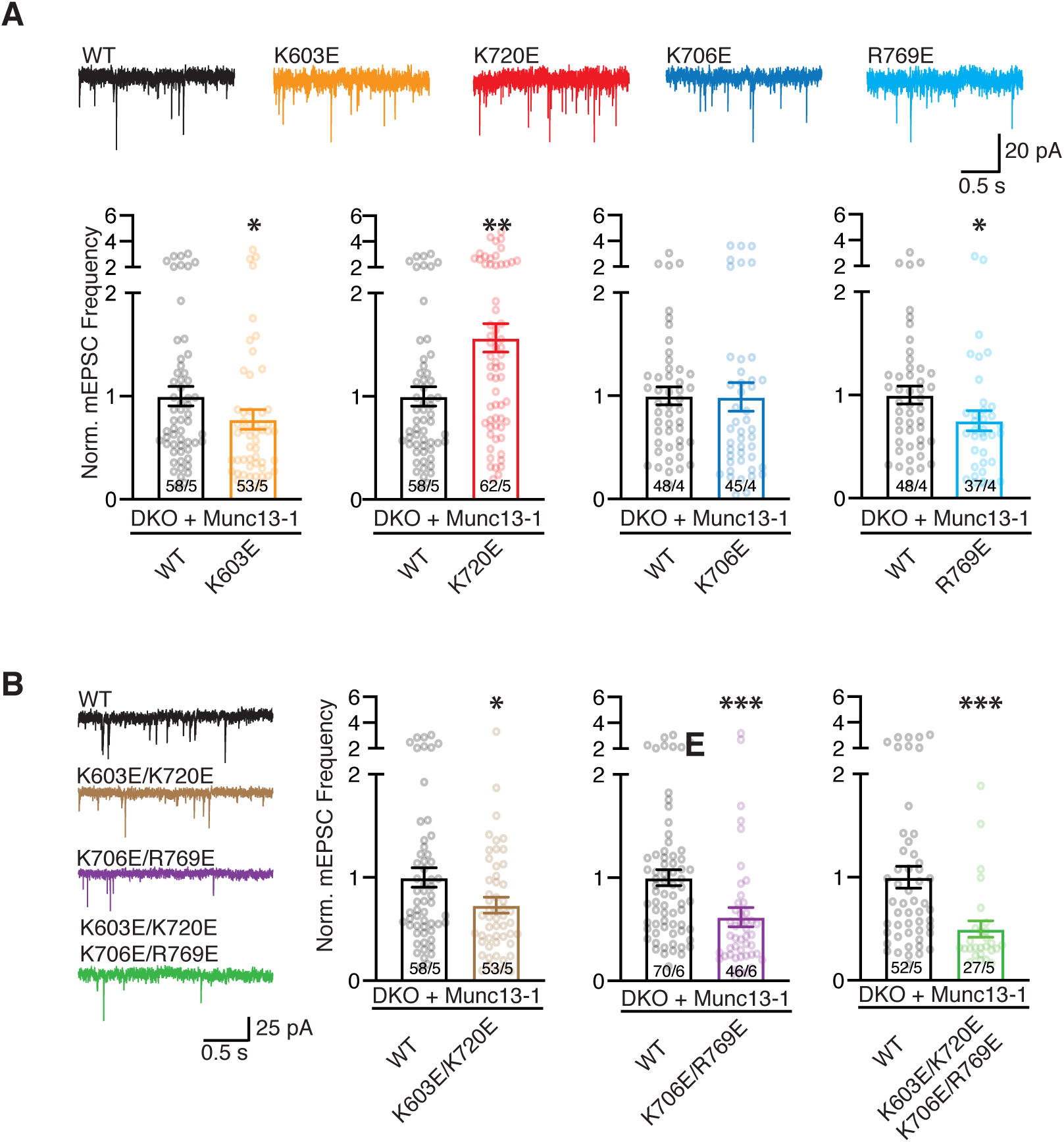
Quantification of spontaneous neurotransmitter release of point mutations within the Munc13-1 C_1_-C_2_B region. **A.** Example traces of mEPSCs and bar plot of mean frequencies normalized to the corresponding Munc13-1 WT (black) from Munc13-1 polybasic mutants: K603E (orange), K720E (red), K706E (blue) and R769 (light blue). **B.** Example traces of mEPSCs and bar plot of normalized mean frequencies from the double and quadruple mutations introduced in the C1-C2B region. Circles in bar plot represented normalized values per neuron. Numbers in bars corresponded to the cell number/culture number. Significances and p-values were determined using the non-parametric Mann-Whitney U test. *P<0.05; **P<0.01, ***P<0.001.

**Figure 8-Figure supplement 1.**
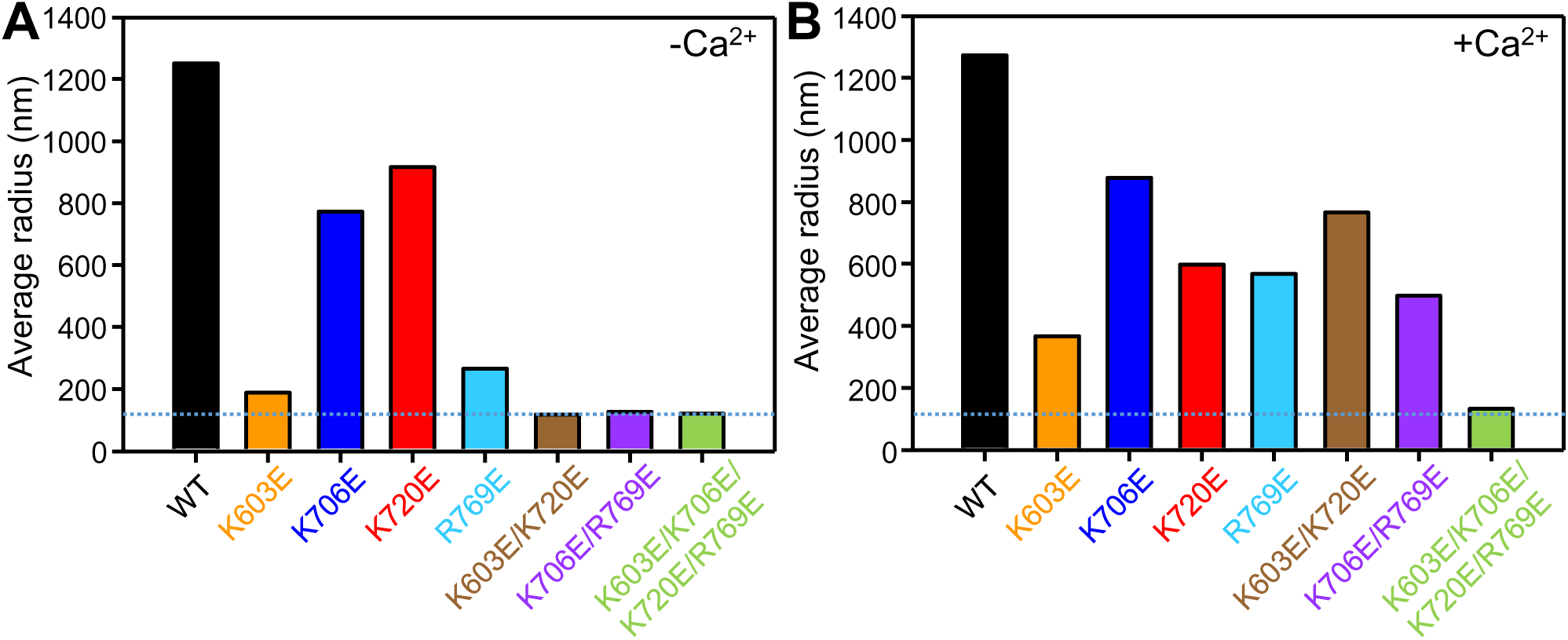
Semi-quantitative analysis of the liposome clustering assays. **A,B.** The particle size distributions observed in the liposome clustering assays after addition of the corresponding C_1_C_2_BMUNC_2_C fragment in the presence of 0.1 mM EGTA (**A**), and after addition of 0.6 mM Ca^2+^ to the same sample (**B**) (shown in Fig. 7) were converted to average radii by calculating a populating weighted average, i.e. the sum of terms obtained by multiplying the radius of each bin by the corresponding population. Note that the equations used to derive the populations of each particle size bin break down for large radii. Therefore, these data need to be interpreted with caution, and the difference between the average radii calculated for the K720E C_1_C_2_BMUNC_2_C mutant in the absence and presence of Ca^2+^ cannot be considered meaningful.

**Figure 10-Figure supplement 1.**
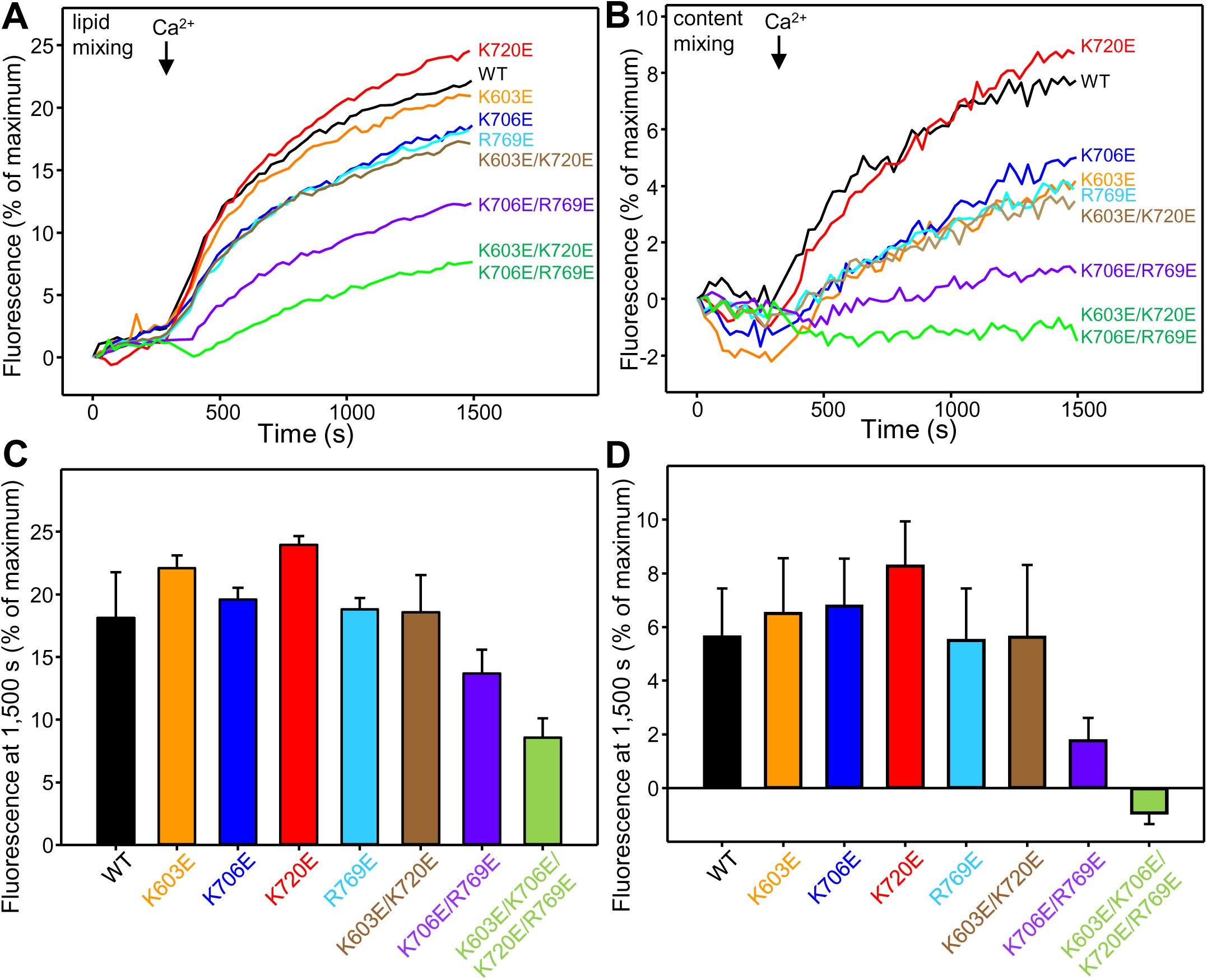
Mutations in basic residues of the C_1_-C_2_B region differentially impair the ability of the Munc13-1 C_1_C_2_BMUNC_2_C fragment to stimulate liposome fusion. **A,B.** Lipid mixing (left) between VSyt1- and T-liposomes was measured from the fluorescence de-quenching of Marina Blue-labeled lipids and content mixing (right) was monitored from the development of FRET between PhycoE-Biotin trapped in the T-liposomes and Cy5-Streptavidin trapped in the VSyt1-liposomes. The assays were performed under analogous conditions as those used for Fig. 9A,B, but the VSyt1 liposomes contained a synaptobrevin-to-lipid ratio of 1:10,000 and a synaptotagmin-1-to-lipid ratio of 1:1,000, and the T-liposomes contained a syntaxin-1-to-lipid ratio of 1:5,000 and a dodecylated SNAP-25-to-lipid ratio of 1:800. Experiments were started in the presence of 100 μM EGTA and 5 mM streptavidin, and Ca^2+^ (600 μM) was added after 300 s. **C,D.** Quantification of the lipid mixing (**C**) and content mixing (**D**) observed after 1,500 s in reconstitution assays performed under the conditions of Panels **A,B**. In **C,D**, bars represent averages of the normalized fluorescence observed in experiments performed at least in triplicate, and error bars represent standard deviations.

